# Mutational landscape and molecular bases of echinocandin resistance in *Saccharomyces cerevisiae*

**DOI:** 10.1101/2024.07.21.604487

**Authors:** Romain Durand, Alexandre G. Torbey, Mathieu Giguere, Alicia Pageau, Alexandre K. Dubé, Patrick Lagüe, Christian R. Landry

## Abstract

One of the front-line drug classes used to treat invasive fungal infections is echinocandins, which target the fungal-specific beta-glucan synthase (Fks). Treatment failure due to resistance often coincides with mutations in three protein regions defined as hotspots. Unfortunately, the scarcity of the mutational data reported, combined with the large size and membrane-embedded nature of the enzyme hinder any effort to characterize genotype-phenotype links. Recent advances in solving the structure of Fks bring us one step closer to reliable predictions of the binding modes of each echinocandin. To help with that endeavor, we used molecular dynamics simulations to develop a membrane-embedded model of Fks that captures key structural and environmental features. Our results show that the three hotspots shape a single solvent-exposed binding cavity, hinting at the orientation and positioning of echinocandins. This structural framework is integrated with deep-mutational scanning to comprehensively assess the impact of mutations across the three hotspots in the model yeast *Saccharomyces cerevisiae*. We elucidate several key molecular bases of resistance to the three most widely used echinocandins; anidulafungin, caspofungin and micafungin and provide clues to better understand intrinsic resistance of critical fungal pathogens.

**One sentence summary:** Key residues at specific positions in Fks hotspots lead to echinocandin-specific resistance.

## Introduction

Echinocandins are among the most important compounds used to combat human fungal infections. They are routinely administered to treat invasive candidiasis and often represent the only therapeutic option against the emerging pathogen *Candidozyma auris* (Pappas et al. 2016; Chowdhary et al. 2017). Unfortunately, their efficacy is increasingly compromised by the acquisition of resistance, which most often arises from mutations in the drug target: the β-1,3-glucan synthase Fks1 (Lee et al. 2023). Mutations identified in clinical isolates with minimum inhibitory concentrations higher than clinical break points, including strains isolated upon treatment failure, are concentrated in three regions of the coding sequence, which have hence been referred to as “hotspots” (Walker et al. 2010; Johnson et al. 2011; Pfaller et al. 2019). Despite the clinical relevance of echinocandins and the growing threat of resistant pathogens, little is known regarding the exact molecular mechanisms at play or even the nature and number of mutations in the hotspots that can cause resistance.

Echinocandins act as non-competitive inhibitors of the enzyme, although their mechanism of action is still not entirely clear (Garcia-Effron et al. 2009; Garcia-Effron 2020). The correlation between treatment failure and amino acid substitutions within the hotspots has initially led to the hypothesis that echinocandins directly bind these regions (Douglas 2001). Since then, many studies confirmed that there is a correlation between mutations in the hotspots and echinocandin resistance (Bédard et al. 2025). In a pivotal study, Johnson et al. (Johnson and Edlind 2012) significantly improved our understanding of the mechanisms underlying echinocandin resistance. They proposed that all three hotspots are in close proximity to each other and are located close to, or embedded in the outer membrane. They further suggested that echinocandins interact with hotspot 1 and 2 via their peptidomimetic macrocycle, and with hotspot 3 via their lipophilic tail (Johnson and Edlind 2012). In the past two years, the structures of Fks1 from the budding yeast *Saccharomyces cerevisiae*, alone and in complex with the regulatory GTPase Rho1, have been solved by three independent groups by cryo-electron microscopy (Hu et al. 2023; Zhao et al. 2023; Li et al. 2025). Structural models combined with molecular docking analyses supported the energetic favorability of echinocandin binding at the spatially clustered hotspot regions (Jospe-Kaufman et al. 2024). Although these recent advances have not yet fully elucidated the molecular mechanisms of resistance, they point to the hotspot regions as key determinants of echinocandin recognition and susceptibility.

A major concern with the emergence of resistance mutations in fungi is the extent of cross-resistance, especially given the limited number of antifungal drug classes and the small number of available compounds within each class (Puumala et al. 2024). If a given mutation conferred resistance to only a single echinocandin, diagnostic testing could guide clinicians toward an alternative molecule within the same class rather than discontinuing echinocandin therapy altogether. To comprehensively assess the extent of cross-resistance, scarce clinical data are insufficient (Bédard et al. 2025). Instead, screening exhaustive libraries of mutants generated in the laboratory provides the only viable approach to quickly and precisely map the full landscape of echinocandin resistance mutations. A notable example of reported mutant conferring echinocandin-specific resistance is the caspofungin-specific resistant W695C substitution in hotspot 3 (Johnson et al. 2011). If this trend were confirmed, it would support a model in which echinocandins bind directly to the hotspot region but adopt distinct binding orientations depending on the compound. However, how such large molecules can engage hotspots located deep within the membrane remains unclear. Further experimental and computational studies are required to better understand how echinocandins interact with these membrane-embedded regions. The presence of polar amino acids within these hotspots may create locally favorable microenvironments that accommodate the amphipathic nature of echinocandins. Clarifying this apparent discrepancy should provide valuable clues regarding their positioning and orientation.

In this study, we combined explicit-solvent molecular dynamics (MD) simulations with deep-mutational scanning (DMS) to investigate the structural and functional bases of echinocandin resistance. MD simulations based on the recently solved structure of Fks1 were used to characterize solvent accessibility and physicochemical features of the hotspot region. In parallel, DMS was used to generate and functionally profile all possible single amino acid substitutions across the three resistance hotspots of Fks1. Mutant libraries were screened against three clinically relevant echinocandins: anidulafungin, caspofungin and micafungin to assess resistance and cross-resistance profiles. Together, these approaches provide mechanistic insights into how specific mutations differentially affect echinocandin susceptibility and help explain intrinsic resistance observed in certain fungal pathogens.

## Results

### Water-filled hotspot cavity features shaping echinocandin susceptibility

Although cryo-EM has resolved the transmembrane helices of Fks1 at high resolution, these structures provide limited information on the organization of the surrounding membrane.

Previous mutational studies, together with structural and modeling analyses, suggest that the three canonical resistance hotspots collectively define a putative echinocandin binding cavity (Johnson and Edlind 2012; Hu et al. 2023; Jospe-Kaufman et al. 2024). One such study used a molecular docking approach to predict the binding mode of anidulafungin and the structurally close rezafungin (Jospe-Kaufman et al. 2024). Given the complexity of the system, different approaches are warranted to gain insight into the biochemical nature of this putative binding pocket. Structural inspection reveals several polar residues embedded within these membrane-facing regions.

Because membrane plasticity can modulate solvent accessibility and ligand accommodation, we performed three independently constructed 1-μs MD trajectories of Fks1 embedded in a lipid bilayer and solvated with explicit water molecules and neutralizing ions. For each system, only the final 300 ns were analyzed to sample water distribution within the hotspot region. Water-density maps derived from these trajectories revealed a water-filled pocket between the helices of hotspot 1 and hotspot 2, extending downward toward the base of the helix at hotspot 3 (residue D691) (Fig. 1 and S1 Fig). This water pocket is stabilized by several polar residues within the hotspot cavity, including K632, E635, S636 (adjacent to HS1), D684 and D691 (HS3), as well as R1357 and S1361 (within and adjacent to HS2, respectively). These results indicate that the upper portion of the hotspot cavity is accessible to extracellular solvent, whereas the lower portion remains embedded within the lipid bilayer. This configuration supports a model in which echinocandin binding takes place in a partially hydrated pocket, where solvent accessibility may modulate both ligand orientation and the balance between polar and hydrophobic interactions. These findings align with the model proposed by Johnson and Edlind (Johnson and Edlind 2012), and highlight the crucial role of the echinocandins’ hydrophobic tail in ensuring stable binding and proper orientation within the cavity. This hydrophobic moiety is critical for inhibition, as first demonstrated with cilofungin, the first semi-synthetic echinocandin drug developed (Taft and Selitrennikoff 1990).

**Fig. 1.**
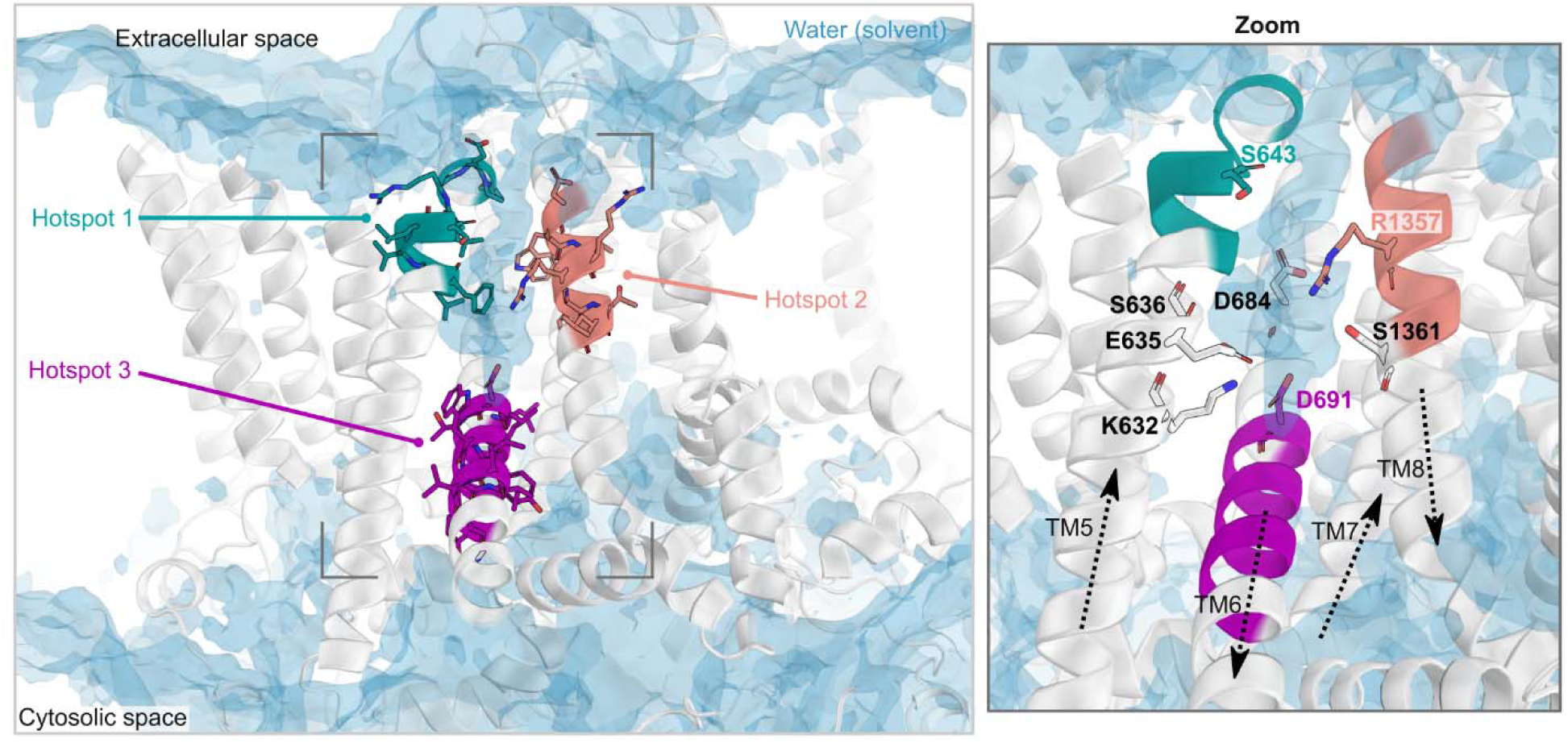
A membrane-embedded water pocket connecting Fks1 hotspots defines the structural bases for echinocandin binding. A water occupancy map (blue surface, isocontour 0.2), computed from three independently constructed MD trajectories of Fks1 (white ribbon) in the absence of echinocandin revealed a water-filled pocket in the transmembrane region. This feature was located primarily between hotspot 1 (dark cyan; F639 to P647) and hotspot 2 (salmon; D1353 to L1360), extending toward hotspot 3 (purple; L690 to N700). The dark gray corners outline the region shown in the magnified view on the right, where only polar residues contributing to the formation of the water pocket are highlighted. Transmembrane helices TM5-8 are shown, with arrow directions indicating the N- to C-terminal orientation of the protein sequence. In this magnified view, the structure was slightly tilted to better visualize these features. Figures were generated using PyMOL v2.5.5 (Schrödinger, LLC 2015).

While this model successfully captures key determinants of echinocandin recognition, it does not fully account for the physicochemical complexity of the binding site revealed by our simulations. In particular, the coexistence of the water-filled pocket and surrounding lipids creates a heterogeneous environment that cannot be reproduced by conventional implicit-solvent docking approaches. To test how this heterogeneous cavity contributes to echinocandin binding and resistance, we generated 13 Fks1 single mutants. These included residues from hotspot 1 and 3, a subset of the polar residues lining the water-filled pocket identified in our MD analysis (E635, D684, and D691), as well as several residues positioned around the cavity that appeared potentially important for ligand binding (F639, R645, R1332, K1334, and K1336).

As expected, the hotspot 3 mutant D691V conferred resistance to all three echinocandins (S2 Fig). The peptidomimetic core of echinocandins is likely positioned within the water-filled pocket, where it may form polar interactions with surrounding residues. Substituting D691 with a nonpolar valine could disrupt these polar interactions, thereby weakening ligand binding across all compounds and contributing to pan-echinocandin resistance.

Similarly, substitution of E635 with either a serine or an arginine resulted in pan-resistance (S2 Fig), consistent with the idea that altering charge at this position may perturb the local hydration network and affect protein-ligand interactions. In contrast, replacing E635 with a valine conferred resistance only to anidulafungin and micafungin, but not caspofungin. This substitution removes hydrogen-bonding capacity and introduces a hydrophobic side chain, which could differentially affect interactions with echinocandin-specific functional groups.

At position D684, substitution with arginine conferred pan-echinocandin resistance, whereas valine or serine did not. This pattern suggests that both charge and the geometry of interactions at this site may contribute to ligand susceptibility. Altogether, these results point to the importance of residues shaping the membrane-solvent interface in modulating echinocandin binding.

By contrast, mutations at R1332, K1334 or K1336, whether to hydrophobic or negatively charged residues, did not confer resistance to any echinocandin (S2 Fig). However, nearly all substitutions at these positions resulted in substantial fitness cost in the absence of drug (S2 Fig), suggesting that these residues contribute to protein function. The EL4 loop, which contains these residues, is defined by two highly conserved cysteines (C1328 and C1345) forming a disulfide bridge. We retrieved 216 unique EL4 sequences from the multiple sequence alignment of 404 Fks homologs. The cystein pair was present in all loops, with 187 (87%) containing at least one lysine or arginine. EL4 was also recently shown to be essential for enzymatic activity (Zhao et al. 2023). Given its flexibility and apparent redundancy among positively charged residues, this loop may tolerate substitutions better than critical polar residues such as E635 or E691.

### Deep-mutational scanning of Fks hotspots

In order to enhance our understanding of the contribution of hotspot mutations to resistance and cross-resistance, we performed deep-mutational scanning of all three hotspots of Fks1 by engineering the corresponding positions in the genome of *S. cerevisiae*. In this organism and many pathogenic fungi such as the closely related yeast *Nakaseomyces glabratus*, Fks1 homologs and particularly the three hotspots are very conserved. Specifically, 24 out of 28 residues across the three hotspots of Fks1 in *S. cerevisiae*, *N. glabratus*, *C. auris* and *Candida albicans* are identical, with the predicted structures for these homologs aligning with a RMSD of 3.06 □ (Fig. 2A and S3 Fig). *S. cerevisiae* is not only the most appropriate model to generate extensive mutagenesis data directly on the protein that was used for solving the structure, the conservation of these regions may reflect conservation of the molecular mechanisms across species.

**Fig. 2.**
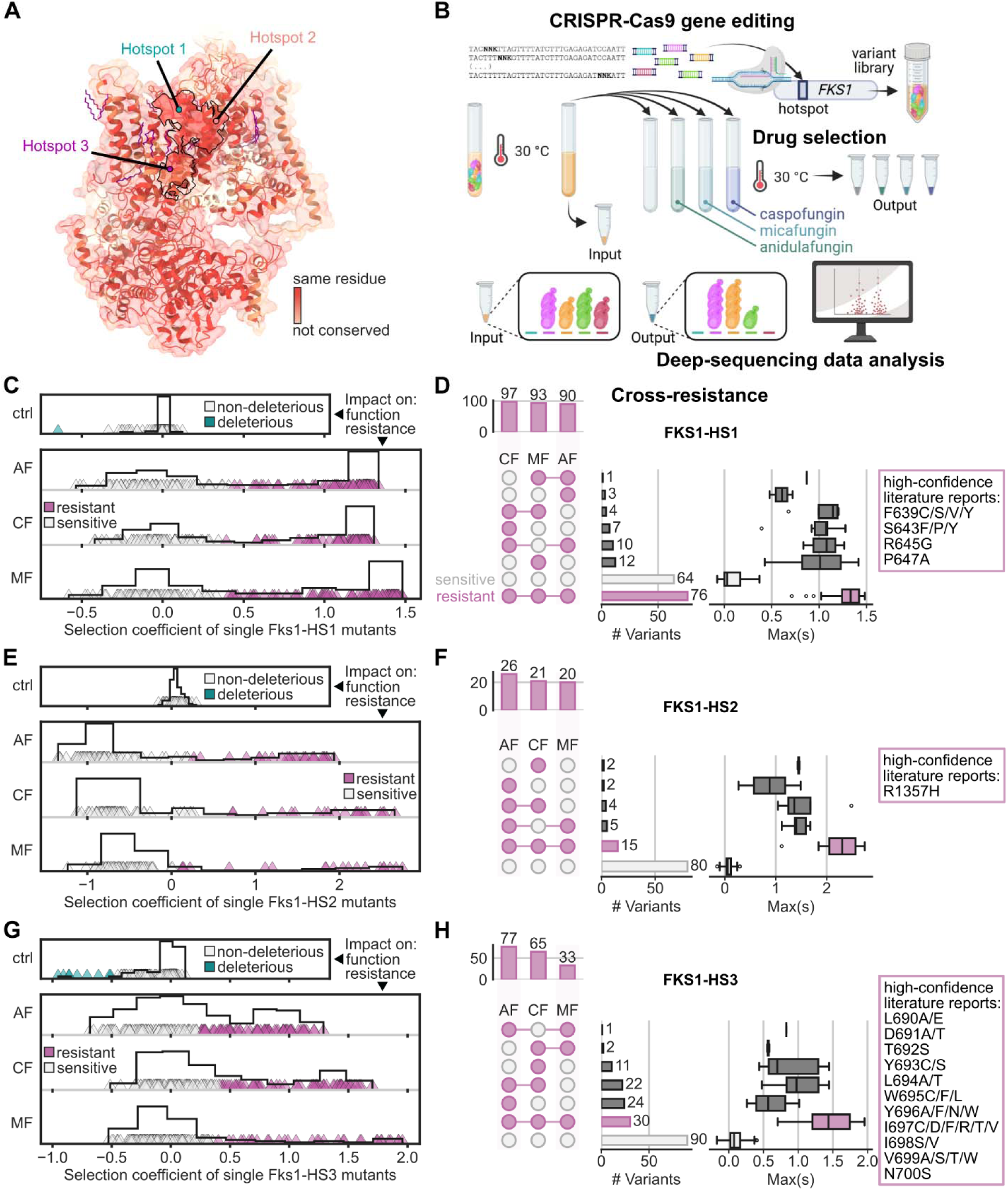
Deep-mutational scanning of β-glucan synthase (Fks) hotspots identifies specific and cross-resistance mutations to echinocandins. **(A)** Fks1 structure from *S. cerevisiae* (PDB: 7XE4) (Hu et al. 2023), visualized using UCSF ChimeraX v1.8 (Pettersen et al. 2021), colored according to Jalview’s conservation score (Livingstone and Barton 1993) from the structural alignment of Fks1 from *N. glabratus*, *Candida albicans* and *C. auris* (AlphaFold predictions for these three homologs, as shown in S3B Fig). (**B**) Simplified experimental design for the DMS assay, created with BioRender (https://BioRender.com/gorwv4e), showing how a library of single mutants can be obtained by integrating oligonucleotides with NNK codons at the endogenous locus of any *FKS* hotspot. The library is cultured and subjected to bulk competition. Following genomic DNA extraction and amplicon sequencing, selection coefficients are calculated from the change in allele frequencies before/after screening. (**C, E, G**) Distribution of effects for single mutants of (**C**) hotspot 1 (**E**) hotspot 2 and (**G**) hotspot 3 of Fks1 (including wild-type) in each condition. In the control condition (ctrl), the effect relates to protein function, whereas in the presence of drug (AF, anidulafungin; CF, caspofungin; MF, micafungin), the effect informs on resistance. Effects were quantified using the selection coefficient (the wild-type is expected to have a selection coefficient around 0). Variants were classified based on their effect using a Gaussian mixture model (see methods). The black line indicates the distribution regardless of the classification. (**D, F, H**) UpSet plots showing the number of unique (**D**) Fks1-HS1, (**F**) Fks1-HS2 and (**H**) Fks1-HS3 amino acid sequences classified as resistant (pink circles) or sensitive (gray circles) to each drug, and their overlap. For example, in panel D, 76 single mutants are resistant to all three echinocandins. Barplots at the top left of each panel indicate how many variants are resistant to the indicated drug. On the right, a boxplot shows the corresponding maximum selection coefficient across drugs for each set. The mutations listed are high-confidence reports from previously published studies for which there was at least one observation of resistance that matched the results presented here (S1-S2 Data). These reports include observations made from several fungal pathogens featured on the WHO priority list (WHO fungal priority pathogens list to guide research, development and public health action 2022).

Mutant oligo libraries (NNK codon at each position) were amplified and integrated at the endogenous hotspot loci using CRISPR-*Cas9* editing (Fig. 2B). We also amplified libraries containing unique hotspot sequences found in homologous *FKS* genes to study the link between hotspot sequence content and intrinsic resistance. The mutant strains were cultured overnight without selective pressure. They were then exposed to the three echinocandins individually (anidulafungin, caspofungin, micafungin). The concentrations used were estimated to inhibit around 80% of wild-type growth in most conditions. Cells were harvested before and after two consecutive rounds of selection and the genomic loci corresponding to the three hotspots were deep sequenced to estimate the extent of selection acting on each variant using changes in relative frequencies.

For all mutated hotspots, at least one nucleotide variant could be successfully sequenced in the initial pools for every expected amino acid variant for both types of libraries (NNK and homologous hotspots). However, sequencing the diversity in the DMS input revealed that culturing overnight in rich medium was sufficient for several variants to be depleted, most likely because of their impact on fitness in the absence of drug (or the impact of surviving variants, S4 Fig) and/or due to experimental bottleneck(s). Depletion was not systematic (S5 Fig), ruling out experimental bottleneck(s) as possible cause. Instead, while hotspot 1 showed little to no depletion (at least 95% of variants remaining, S5 Fig), as few as 63% of single mutants of hotspot 2 remained after culturing overnight. An intermediate trend could be observed for hotspot 3 (81% of variants remaining, S5 Fig). The average mutant coverage at T0 for the corresponding sequencing run exceeded 700X, which leads us to infer that the depleted variants had a deleterious effect on fitness.

Selection coefficients were computed for each nucleotide variant and then averaged within each amino acid variant. Replicates among cultures correlated well, with a Spearman’s correlation coefficient ranging from 0.60 to 0.98 for drug conditions (S6 Fig).

Overall, we obtained a bimodal distribution of effects when selecting for echinocandin-resistant mutants, regardless of the drug used (Fig. 2C). This bimodality is consistent with strong selective pressure allowing only resistant mutants to grow significantly. In the case of hotspot 2 and hotspot 3 mutants, the two modes were less clearly defined, mostly because there were more sensitive mutants than for hotspot 1 (Fig. 2). We also observed an average selection coefficient for sensitive mutants in hotspots 2 and 3 below 0 (Figs. 2E,G and S7 Fig), again indicative of fitness cost. In the case of hotspot 2, harsh selection manifested from the generation of libraries (see methods) up until the end of the first screening round (t1), with cultures growing systematically slower than other conditions, reaching fewer mitotic generations (S4B Fig).

Because of survivorship bias, there were no hotspot 2 mutants classified as deleterious in the absence of drug (Fig. 2E). In contrast, fitness costs associated with mutating hotspot 3 did not manifest until the screening round, which led to several mutants being classified as deleterious (Fig. 2G). Finally, selection coefficients for hotspot 2 mutants were also biased by the low number of silent mutants that could be engineered using NNK libraries given the wild-type sequence, resulting in a shifted distribution of coefficients. Despite these constraints, we were still able to distinguish sensitive from resistant mutants for all three hotspots.

### Classification and validation of mutational effects

A total of 465 Fks1 single mutants were confidently classified as either sensitive or resistant to caspofungin, micafungin or anidulafungin across hotspots 1, 2 and 3, using a Gaussian mixture model (Fig. 2 and S7 Fig).

In order to validate the effects of these mutations on resistance, 28 single mutants in the hotspot 1 of Fks1 spanning the full fitness range were recreated individually by genome editing. The estimated fitness values strongly correlated with the selection coefficients from the bulk-competition assay (Spearman’s *r* = 0.92-0.94 across drugs, *p*-value <= 1.1e-10, S8 Fig).

We find that 61 mutants across all three hotspots (13% of classified mutants) were specifically resistant to a single echinocandin molecule (Figs. 2D,F,H). 50% were resistant to at least one echinocandin drug, while 26% were resistant to all three tested molecules. More than half of these pan-echinocandin resistant mutants were in hotspot 1 (n=76, Fig. 2D), which may partly be explained by the lower fitness cost compared to the two other hotspots. On the other hand, resistance in hotspot 1 does not seem directed to any specific echinocandin. This contrasts what we observe with hotspot 2 resistant mutants, almost all of which confer resistance to anidulafungin (Fig. 2F). This pattern, along with previous observations (Jospe-Kaufman et al. 2024), suggests that anidulafungin may occupy the mixed hydrophobic-polar pocket involving W1354 and R1357. Finally, many hotspot 3 mutants were resistant to both anidulafungin and caspofungin (Fig. 2H), potentially indicating a role played by these residues in drug binding.

Several hotspot mutations have previously been engineered or identified in lab-evolved and clinical strains of a number of fungal species, including seven that feature on the WHO fungal priority pathogens list (WHO fungal priority pathogens list to guide research, development and public health action 2022): *C. auris*, *Aspergillus fumigatus*, *C. albicans*, *N. glabratus*, *Candida tropicalis*, *Candida parapsilosis* and *Pichia kudriavzevii* (S1 Data, obtained by parsing the FungAMR database (Bédard et al. 2025)). Our results validate the reported resistance (or sensitivity) phenotype for 43 of these mutants (S2 Data). As pointed out in the FungAMR study, which aggregates most reports of antifungal resistance to date, resistance among species is likely to be acquired through parallel evolution to the same variants (Bédard et al. 2025). Our results support this hypothesis, suggesting that observations made in one species are informative for predicting resistance phenotypes in other species.

### Echinocandin-specific resistance depends on the position and electrostatic profile of substituted residues

Next, we used the full deep-mutational scanning dataset to rationalize the molecular bases of resistance. Our data revealed a strong position-dependent pattern of echinocandin resistance (Fig. 3B-D). Nearly all mutations at positions 639, 643, 646 and 647 conferred resistance to anidulafungin and caspofungin, while resistance to micafungin was primarily associated with mutations at positions 639, 643, 645 and 647 (Fig. 3D). Notably, these resistance-conferring mutations imparted little to no fitness cost (Fig. 3A), which may partly explain why they are so frequently reported in clinically resistant isolates (S1 Data).

**Fig. 3.**
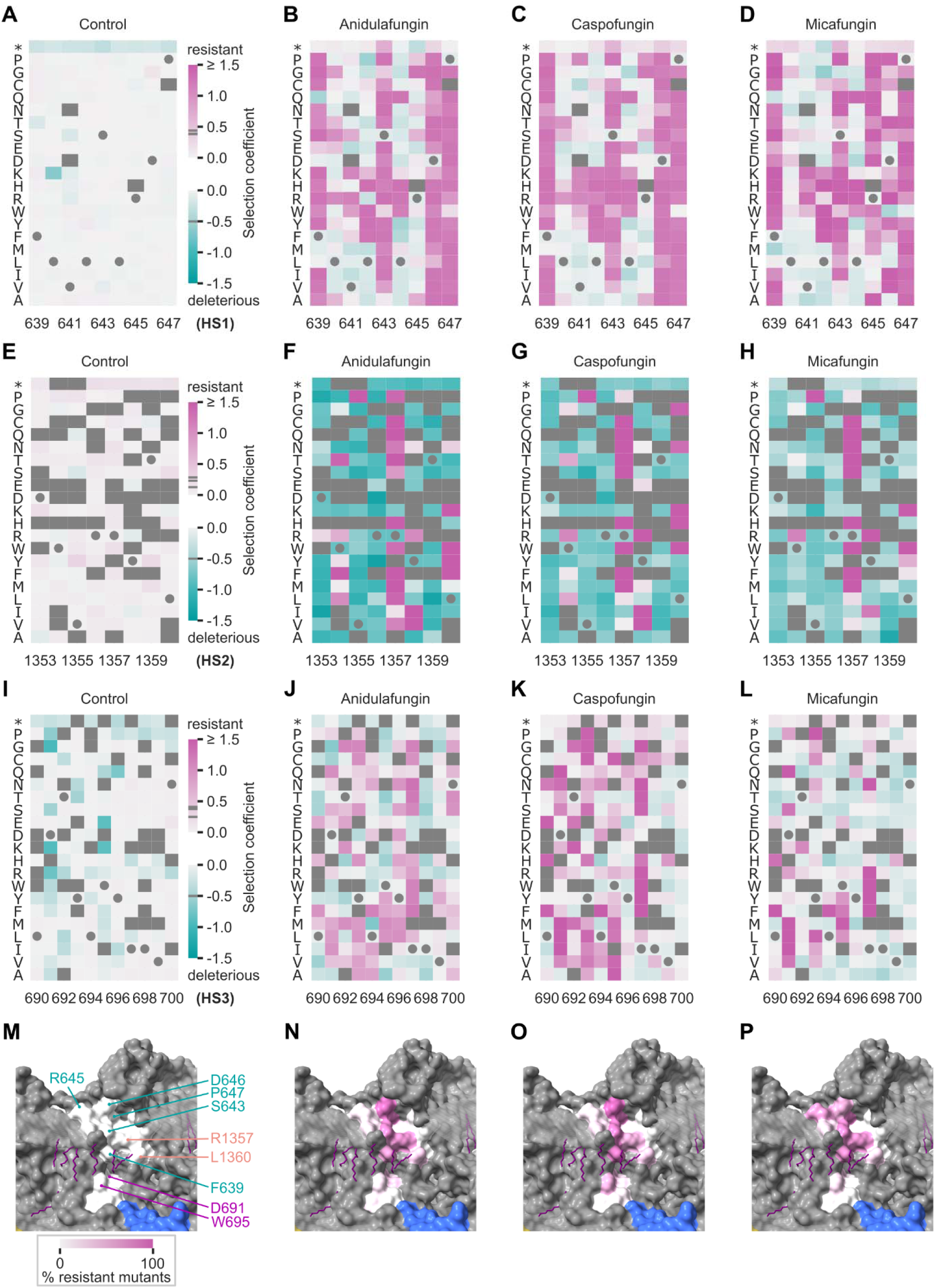
Crucial positions in Fks1 for echinocandin resistance. (**A-L**) Selection coefficients (median of synonymous variants) of individual mutants in hotspot 1 (A-D), hotspot 2 (E-H) and hotspot 3 (I-L) of Fks1 in control (A, E, I), anidulafungin (B, F, J), caspofungin (C, G, K) and micafungin (D, H, L). The wild-type sequence is indicated by gray circles. We computed the selection coefficients of five mutants absent from the initial mutant libraries: F639C, L640D, L642G, P647N and P647Q in validation assays. In all panels, some variants have a selection coefficient above 1.5 but were set to 1.5 to ease the data representation (see the distribution of values in Fig. 2). (**M-P**) For each condition (same order as previous panels), all three hotspots in the structure of Fks1 are colored according to the proportion of mutants classified as resistant at each position (relative to the number of mutants for which we were able to obtain a selection coefficient). The labels of key residues are colored according to the hotspot, residues in blue at the bottom of each panel belong to the catalytic domain, while residues in gray correspond to other non-mutated residues in the transmembrane domains. The approximate location of membrane bilayers is shown by two semi-opaque circular planes (spheres overlapping with the hotspots are hidden for clarity). Lipids present in the structure (Hu et al. 2023) are represented in purple sticks. The corresponding panels were generated in UCSF ChimeraX v1.8 (Pettersen et al. 2021).

Hotspot 1 is located at the water-membrane interface (Fig. 1). F639 is embedded in the hydrophobic core of the membrane, S643 lies near the lipid-water interface, and R645 and D646 are fully solvent-exposed (Fig. 1). In this region, the alpha-helix undergoes a transition in orientation: residues upstream F639 adopt a transmembrane orientation, whereas those following P647 align more parallel to the membrane surface and are positioned toward the extracellular side of the protein. This shift is favored by a combination of structural features, including the helix-breaking effect of P647, the propensity of S643 to induce local distortion in the helix backbone (Deupi et al. 2010), the membrane anchoring of F639, and the strong solvation of R645 and D646. Mutations that disrupt this delicate balance may promote helix extension and alter the binding pocket structure.

Relatively few substitutions at positions 640 and 644 conferred resistance (Fig. 3B-D), consistent with the location of these leucine residues within the hydrophobic pocket. Because they do not directly line the cavity, substitutions at these positions are less likely to perturb echinocandin binding than mutations affecting residues that define the cavity surface. A notable exception is the L644W mutation, which retained hydrophobic character but introduced substantial steric bulk, effectively filling, and potentially occluding the pocket, leading to cross-resistance.

A key distinction across the three drugs is the mutational sensitivity of positions 645 and 646. Substitutions at D646 more frequently conferred resistance to caspofungin and anidulafungin, whereas mutations at R645 were more commonly associated with micafungin-specific resistance.

In the case of hotspot 2, most substitutions associated with resistance were at positions 1357 and 1360 (Fig. 3). However, they were outnumbered by substitutions which were both sensitive and deleterious in the absence of drug. The magnitude of their effect often lying under the quality threshold explains the apparent missing data in the heatmaps and resulted in inflated selection coefficients (with many sensitive mutants appearing as if they were very deleterious). Despite these technical artefacts, it is very clear that most hotspot 2 mutants were at least somewhat deleterious. In particular, mutations introducing negatively charged residues, such as aspartate or glutamate, consistently resulted in a strong fitness cost (Fig. 3E). This sensitivity is likely due to the proximity of hotspot 2 to residues F1311 (TM7b) and F1475 (TM12), which have been proposed to form part of the exit of the putative glucan translocation channel (Hu et al. 2023; Li et al. 2025). Disruption of the delicate balance between TM8, which contains hotspot 2, and the neighboring transmembrane helices TM6, TM7b and TM12 could interfere with gating of the translocation channel at this site. This interpretation is further supported by the observation that mutations at positions 1353, 1356, and 1357 incurred a lower fitness cost (Fig. 3E), likely because these residues are exposed to the solvent or lipid environment rather than being involved in helix-helix packing.

Consistent with our MD-based results, seven missense mutations at position 691 in hotspot 3 (including the recreated mutant D691V) conferred pan-echinocandin resistance (Fig. 3J-L), suggesting a central role of the aspartate in drug binding. Also in hotspot 3, ten missense mutations at position 695 conferred caspofungin resistance, with five of them conferring caspofungin-specific resistance, including W695C as previously reported (Fig. 3J–L) (Johnson et al. 2011). With limited biophysical data available, this phenotype was proposed to result from differential binding of the echinocandin lipid tails. Finally, we note that most hotspot 3 mutants conferring micafungin resistance are pan-echinocandin resistant (Fig. 2H), further highlighting the distinct role that hotspot 3 plays depending on the drug.

### Mutational landscapes of paralog hotspots overlap

In several clinically relevant fungi, as well as *S. cerevisiae*, there are at least two duplicated *FKS* genes (Denning 2003). In our model yeast, *FKS2* is expressed at a very low level during vegetative growth, unless *FKS1* is repressed or inactivated (Mazur et al. 1995). As a result, both paralogs are synthetically lethal, which allows editing them directly in the genome, although our results suggest that many *FKS1* mutations cannot be masked by *FKS2* (for example in hotspot 2). To assess the impact of amino acid substitutions in Fks2 and examine if the patterns of resistance are conserved between these two paralogs, we built libraries for hotspot 1 and hotspot 2 as we did for Fks1 and measured fitness cost with and without repression of *FKS1*, as well as fitness effects in the presence of drug with repression of *FKS1*.

As expected for synthetic lethal gene duplicates, introducing premature stop codons in *FKS2* hotspots had little effect in the WT background but resulted in a strongly deleterious effect when *FKS1* was repressed (S9 Fig). Despite lower resolution for *FKS2* compared to *FKS1*, the pattern observed for other substitutions largely corroborated those obtained with Fks1 (S9 Fig), which we attribute to the high degree of protein sequence identity (87.7%) and functional redundancy between the two paralogs. More specifically, substitutions at positions 662, 1376 and 1379 (homologous to S643, R1357 and L1360 in Fks1, respectively) frequently conferred pan-echinocandin resistance, while substitutions at R664 (homologous to R645 in Fks1) led to micafungin-specific resistance (S9 Fig). Although we did not mutate hotspot 3 in Fks2, previous studies have reported overlaps in resistance profiles for equivalent mutations across both paralogs. For example, the Fks2 W715L mutation (homologous to W695 in Fks1) was shown to confer resistance to anidulafungin and micafungin in *N. glabratus* (Bordallo-Cardona et al. 2018). This suggests that mutations in *FKS2* could also have similar effects on resistance as those in *FKS1*.

To confirm that native expression of *FKS2* has little impact on the mutational landscape observed for Fks1, we also screened Fks1 libraries upon repression of *FKS2*. Since the same libraries were constructed in two genetic backgrounds, this could also be used as an additional measure of replicability. We confirmed that the selection coefficients correlated well between the two libraries and backgrounds, with or without repression of *FKS2* (Spearman’s *r* = 0.83-0.91 across drugs, *p*-value <= 7.0e-33, S10 Fig).

### DMS data of wild-type homologous sequences corroborate previous reports of intrinsic resistance

If hotspot sequences are enough to influence echinocandin resistance, measuring the adaptive power of wild-type homologous hotspot sequences during the same screen as our DMS may provide clues as to why some homologs appear insensitive to echinocandins. To evaluate this, we spiked the initial libraries of single mutants with homologous hotspot sequences. These sequences often differ from the wild-type hotspot found in *S. cerevisiae* by several amino acid residues. Comparing the phenotype conferred by close sequences may help distinguish crucial residues playing a role in intrinsic resistance to echinocandins from those that result from standing genetic diversity. In order to identify associations between specific amino acid residues at key positions and resistance phenotype across all classified hotspots, we generated enrichment logos from the DMS data obtained with each tested echinocandin. The relative occurrence of residues in homologous hotspot sequences was converted to a change in information content (S11 Fig), reflecting both the level of conservation across homologs and the degree of enrichment in resistant or sensitive hotspots. Pan-echinocandin resistant hotspots often had tyrosine, leucine or phenylalanine at positions 639, 640 and 644, respectively. Since we find only F639Y confers resistance in our DMS, the enrichment of the leucine and phenylalanine may be due to a lower mutational tolerance at positions 640 and 644 across homologs. In contrast, hotspots classified as sensitive to all three echinocandins often had tyrosine or proline at positions 639 and 647, respectively (S11 Fig). This confirms that a substitution at those positions is likely to affect resistance across homologs. In hotspots 2 and 3, there was almost no enrichment characteristic of resistance, while a single residue was often enriched at each position of sensitive hotspots. This single residue corresponded to the wild-type residue in *S. cerevisiae* in most cases. This indicates that the high level of conservation observed for hotspots found in *C. albicans*, *C. auris* and *N. glabratus* (Fig 2A) is also true for a much broader range of evolutionary distant species (S3A Fig). Such high conservation may be explained by the important role that hotspots 2 and 3 appear to play in the structure and/or protein function of Fks1, resulting in low mutational tolerance despite a potentially wide array of selective pressures being exerted across many distinct ecological niches. Still, we do observe two notable associations at position 695 in hotspot 3: phenylalanine in anidulafungin-resistant hotspots and tryptophane in sensitive hotspots. This corroborates several of our previous observations, including the association between hotspot 3 and anidulafungin and the important role that W695 appears to play in anidulafungin resistance and possibly binding (Fig 2H and S11 Fig).

Next, we examined specific cases where we hypothesized hotspot 1 content was at least necessary to observe intrinsic resistance in key fungal pathogens. We find that substituting the hotspot 1 of *S. cerevisiae* with its homologous counterpart from the fungal pathogen *C. parapsilosis* leads to pan-echinocandin resistance. This supports the hypothesis that the hotspot 1 sequence, which ends with an alanine instead of a proline because of a naturally-occurring polymorphism, is the main cause for the decreased sensitivity exhibited by several strains of *C. parapsilosis*, *Candida orthopsilosis* and *Candida metapsilosis* (Garcia-Effron et al. 2008). Several species of *Mucorales* encode at least two Fks paralogs, each with a different hotspot 1 sequence (S3 Data). Interestingly, one sequence was classified as pan resistant, while the other was classified as sensitive. The former differs from *S. cerevisiae* by substitutions V641A, L644F and P647A, while the latter differs from *S. cerevisiae* by substitutions V641A, L644F and R645K. Although proper validation in the appropriate background is needed, it would seem that again, the presence of an alanine instead of a proline at position 647 is necessary to observe intrinsic resistance in *Mucorales*. The hotspot 1 sequence found in Fks1 from *Fusarium* was also classified as pan resistant. Differing from its *S. cerevisiae* homologous counterpart by a F639Y substitution (among others), the sequence was previously shown to play a critical role in intrinsic resistance of these pathogenic fungal species (Katiyar and Edlind 2009).

## Discussion

Interpreting the impact of resistance-associated mutations on echinocandin susceptibility, as well as guiding the rational improvement of antifungal therapies, requires a comprehensive understanding of resistance-associated mutations.

For echinocandins, this task is particularly challenging because of their large size and intrinsic flexibility, combined with the even greater size, complexity and membrane-embedded nature of their target, Fks1. As a result, traditional molecular docking approaches remain poorly suited to infer binding modes, and one cannot either leverage commonly used computational tools for predicting mutational effects, such as FoldX (Delgado et al. 2019) or Flex ddG (Barlow et al. 2018). Although recent advances have provided insights into the overall architecture and mechanism of action of the enzyme (Hu et al. 2023; Li et al. 2025), they do not fully explain how echinocandins bind, how this binding is altered by mutations, or how such changes lead to resistance.

To address these questions, we first ran MD simulations to characterize the physicochemical environment surrounding the putative binding site of echinocandins in a realistic membrane context. Next, we used DMS to screen a large number of single amino acid mutants. We show that (1) the three known hotspots of Fks1 and their composition shape a single binding pocket partially exposed to the solvent, a feature likely to influence both drug orientation and interaction patterns, (2) hotspot 1 plays a major role in echinocandin resistance, with many substitutions at key positions associated with pan-echinocandin and echinocandin-specific resistance, (3) hotspots 2 and 3 appear to play an important role in protein structure and/or function, with several residues apparently involved in drug binding.

Previous studies have proposed binding modes for echinocandins, as well as for related Fks inhibitors such as ibrexafungerp, based on molecular docking (Jospe-Kaufman et al. 2024; Kumar et al. 2024). Notably, the anidulafungin poses reported by Jospe-Kaufman et al. (Jospe-Kaufman et al. 2024) assumed the presence of lipids within the binding cavity. In our mutational analysis, mutating D691, E635 and D684, residues that appear occluded by lipids in these docking poses, conferred resistance to all three echinocandins (Fig. S2).These results instead support a model in which these residues make direct contributions to drug recognition, underscoring the limitations of docking approaches that do not explicitly account for membrane plasticity and solvent heterogeneity.

Unlike traditional structure-function studies, our approach enabled the estimation of mutational effects of several hundreds of variants, with endogenous expression levels, allowing direct comparison of resistance landscapes for the three most widely used echinocandins. Consistent with clinical and laboratory data, most resistance-associated mutations that have been identified in Fks1, whether it be from lab-evolved or clinical strains of fungi or here, are localized in hotspot 1 (Fig. 2 and S1 Data). Our results further suggest that a lower fitness cost associated with mutations in this region may confer higher mutational tolerance in the absence of drug (Fig. 2).

Within hotspot 1, our data highlight distinct roles for individual residues. F639, located at the membrane-solvent interface, likely anchors the hotspot 1 helix and contributes to its proper positioning within the binding cavity. Substitutions at this position are therefore expected to affect local structure or accessibility, consistent with their frequent association with pan-echinocandin resistance (S1 Data). Our model also provides clues for the mechanisms underlying echinocandin-specific resistance. For instance, mutations at R645 in hotspot 1 led to two distinct mutational effects: one associated with resistance to anidulafungin and caspofungin, and another specific to micafungin (Fig. 3). Our modeling suggests that R645 contributes to solvation-driven positioning of the helix at the water-membrane interface, in a region whose orientation is stabilized by membrane anchoring and neighboring helix-disrupting residues such as P647 and S643 (Fig. 1). Mutations at R645 may therefore alter cavity geometry or accessibility in a manner that differentially affects echinocandin binding. Notably, micafungin is distinguished from other echinocandins by the presence of a sulfate group, raising the possibility that this moiety engages R645 directly. While this hypothesis remains to be confirmed, it is consistent with prior observations that substitutions at this position behave differently across species, including the R645G mutation, which confers micafungin-specific resistance in *C. albicans* (R647G, (Lackner et al. 2014)) but pan-echinocandin resistance in *K. marxianus* (R657G, (Staab et al. 2014)). Further characterization of this specific residue may help uncover the molecular bases of echinocandin-specific resistance, which could help improve drug stewardship in the long run.

While modification of the existing compounds has generated a diversity of available molecules for this essential class of antifungals (Szymański et al. 2022), there is a knowledge gap when it comes to their exact mechanism of action, particularly how each drug-specific binding mode is affected by amino acid changes in the hotspot sequences. Predictions of the echinocandin susceptibility profile from sequence alone could help prescribe the most relevant echinocandin, increasing the chances of treatment success while helping to slow down the development of resistance. This approach has successfully been applied to the etiologic agents of malaria or tuberculosis (Pines et al. 2020; Murphy et al. 2023), but has yet to be implemented in fungi. Our data and recent work on other antifungal drug targets (Després et al. 2022; Bédard et al. 2023) contribute to this collective effort. Here, we spiked our DMS libraries of single mutants with homologous hotspot sequences and looked for associations between hotspot sequence content and resistance. Such an experimental design means that the rest of the gene stays identical to the *S. cerevisiae* background, which in turns ensures that the phenotype measured is not caused by any interaction involving residues outside of the hotspot. If the resistance phenotype observed in the DMS matches the phenotype of the original species, then the wild-type hotspot sequence introduced may at the very minimum be *sufficient* to confer intrinsic resistance (or decreased sensitivity). On the other hand, such an experimental design cannot show that the hotspot sequence is *necessary* to confer intrinsic resistance. In other words, the model can provide insight into why certain species are intrinsically resistant to echinocandins, if there is a significant chance that resistance or reduced susceptibility is indeed caused by residues altering the binding pocket. Other causes of resistance are not covered by this experiment, for example interactions between alleles in diploid species such as *C. albicans*. Finally, inhibiting Fks1 function might not be enough to translate to organismal sensitivity. The most notorious example of this is *Cryptococcus neoformans*, which displays decreased sensitivity (Denning 2003) even though the Fks1 homolog has been shown to be sensitive to all echinocandins in vitro (Maligie and Selitrennikoff 2005).

Our results have also highlighted residues outside of the hotspots that contribute to drug binding. These residues may have been underreported in earlier resistance studies due to diagnostics biases, such as sequencing limited to the “traditional” hotspot regions. Alternatively, they may be less mutationally tolerant than the residues in hotspot 1, possibly due to essential structural or functional roles within the enzyme. This is supported by the high level of conservation observed in wild-type homologous hotspots (S11 Fig). This insight could be leveraged to guide the rational re-design of existing echinocandins to favor stable interactions with these mutationally fragile residues. Strengthening these interactions could slow the emergence of resistance by imposing a greater fitness cost on potential mutations. However, a major assumption of this theory is that the only way to gain resistance is through single substitutions. Recent findings in *N. glabratus* disprove this assumption: resistance-conferring gene conversion events between the *FKS1* and *FKS2* paralogs were shown to produce a chimeric Fks1 that differs at multiple positions, including in EL4 (Zajac et al. 2025). More specifically, this structural rearrangement was identified in clinical isolates from a patient treated with micafungin, and likely an adaptive response to micafungin selection. This resulting chimeric protein included eight amino acid changes, four within or adjacent to EL4 (at positions equivalent to 1327, 1332 and 1336 in *S. cerevisiae*), one in hotspot 2 (V1355I in *S. cerevisiae*) and three close to the active site of Fks1 (positions 1411, 1417 and 1447 in *S. cerevisiae*). These combined changes correlated with an increase in MIC not only for micafungin, but also for fluconazole and voriconazole (Zajac et al. 2025). The authors speculate that such gene conversion events may offer an evolutionary advantage by distributing the resistance burden across multiple positions, avoiding the high fitness cost of single critical mutations, as we’ve observed when mutating R1332, K1334 and K1336 (S2 Fig). To use the popular analogy of the adaptive fitness landscape, there might be a hiking trail around the fitness peak that is EL4.

In summary, by integrating molecular simulations in an explicit membrane environment with large-scale mutational profiling, our study advances the understanding of the mechanisms of resistance of echinocandins. Our results identify hotspot 1 as the primary determinant of resistance, clarify the structural basis of compound-specific effects, and highlight evolutionary constraints shaping resistance pathways. Together, these findings provide a framework for interpreting resistance mutations and for guiding the development of next-generation antifungal strategies.

## Materials and Methods

### Strains, plasmids, primers and culture media

Strains, plasmids, primers and oPools used in this study are listed in S4 Data. The following culture media were used: either YPD (1% yeast extract, 2% bio-tryptone, 2% glucose, with or without 2% agar) or SC (0.174% yeast nitrogen base without amino acids, 2% glucose, 0.5% ammonium sulfate, standard drop-out mix). Most chemical products were acquired from Fisher Scientific or BioShop Canada. When indicated, the following compounds were added to the medium: caspofungin (Cedarlane Labs), micafungin (Toronto Research Chemicals), anidulafungin (Millipore Sigma), hygromycin B (BioShop Canada), G418 (BioShop Canada).

### Genetic constructions

Unless specified, yeast strain construction was done by standard transformation from competent cells and colony PCR validation by standard lithium acetate DNA extraction (Lõoke et al. 2011). First, each hotspot (FKS1-HS1, F639-P647; FKS1-HS2, D1353-L1360; FKS2-HS1, L659-P666; FKS2-HS2, D1372-L1379) was replaced by a hygromycin (HYG) resistance marker in BY4741 (s_001 in S4 Data) and R1158 (s_002) using primers o_001-008 to generate strains s_003-010. In the resulting R1158 strains, the promoter of *FKS1* or *FKS2* was replaced by a doxycycline-repressible promoter using primers o_014-021 to generate strains s_011-014 (Mnaimneh et al. 2004). Strains s_003-010 were used to introduce amplified mutated hotspot sequences: either synthetic libraries (for the deep-mutational scans) or amplicons obtained by fusion PCR (for individual reconstruction of single mutants). In order to reconstruct individual *FKS1*-HS1 mutants, we used primers mainly designed using NEBaseChanger (o_036-092) to amplify two non-overlapping fragments for each mutation. These fragments were individually amplified from genomic DNA of BY4741 using the KAPA HiFi HotStart DNA polymerase (Roche). For each mutation, we diluted and pooled both amplified fragments 1:20 and used 0.75 μL of the obtained pool as template for the fusion PCR using the Q5 High-Fidelity DNA polymerase (NEB). Strain s_003 was transformed as described elsewhere (Ryan et al. 2016). Each reaction was prepared with 20 μL competent cells, 20 μg carrier DNA, 250 ng pCas plasmid (with a guide RNA targeting the hygromycin resistance marker and carrying a G418 (KAN) selection marker), 8 μL PCR-amplified oPool and 200 μL PLATE buffer (PEG 3350 40%, 100 mM lithium acetate, 10 mM Tris-Cl, 1 mM EDTA pH 8.0). Briefly, transformants (selected on YPD + G418) were tested by colony PCR and cultured in YPD to cure the pCas plasmid. We used spot assays to confirm the antibiotic susceptibility profile, ensuring that the constructed strains had lost both selection markers (HYG (hotspot 1) + KAN (pCas)) and Sanger sequenced the mutated region with primer pair o_032-033 to confirm the introduced mutation. The wild-type hotspot was also reconstituted to ensure that the fitness of the obtained strain would not differ from the parental strain in subsequent growth assays.

In addition to missense mutations, we also attempted to introduce all possible codon deletions in the hotspot 1 of *FKS1*, as this type of mutation has previously been reported in *N. glabratus (Garcia-Effron Guillermo et al. 2009; Alexander et al. 2013; Pham et al. 2014; Pfaller et al. 2019)*. However, all attempts yielded no colonies, hinting at a strong fitness cost for these mutations.

We followed essentially the same steps to reconstruct the other individual mutants in Fks1, including N470R, D691V in hotspot 3 (out of three mutations attempted at that position), three mutants at position E635, three mutants at position E684, two mutants at position R1332, two mutants at position K1334 and two mutants at position K1336. The notable difference with hotspot 1 mutants is that we first had to independently replace four more regions in the gene (corresponding to M465-F474, E635-P647, D684-N700 and D1331-L1340 in the protein sequence) with HYG, using primers o_093-099 to generate strains s_043-46. Fragments were amplified with primers o_101-126. The mutated regions were ultimately Sanger sequenced with primers o_032/034/100/. For each deletion strain (s_043-46), the wild-type was reconstituted and the resulting strains were included as controls in the growth assay.

Out of all missense mutations that we tried to engineer, only a few could not be obtained, including P647A, D691S and D691R.

### Deep-mutational scanning (DMS)

#### Design of synthetic libraries

Two types of synthetic libraries for each of five Fks hotspots (FKS1-HS1, FKS1-HS2, FKS1-HS3, FKS2-HS1 and FKS2-HS2) were designed: the first one containing a single NNK codon for each position of the hotspot, the second containing unique codon-optimized sequences encoding for hotspots found in orthologs retrieved from Metaphors (S3 Data) (Pryszcz et al. 2011). To retrieve as many Fks1 and Fks2 orthologous sequences as possible, we used Fks1 from *Aspergillus niger* (M!42720474_ASPNG) and Fks2 from *N. glabratus* (M!115416204_CANGB) as queries. Because of the high percentage of sequence similarity between paralogs, hits for Fks1 sometimes included both paralogs of a given species (e.g. *S. cerevisiae* or *N. glabratus*). In contrast, using Fks2 from *N. glabratus* as the query did not return the paralog sequence, but returned two paralogs from *C. auris*. For this reason, we refer to the corresponding oPools and their associated variants throughout the text as homologous hotspots. Fks1 orthologous sequences were filtered based on their length using the following criteria: minimum of 1,500 residues (80% length of Fks1 from *S. cerevisiae*) and maximum 1,970 residues (105% length of Fks1 from *S. cerevisiae*). Fks2 orthologous sequences were filtered using the same criteria (80% = 1,517 residues, 105% = 1,991 residues). Three sequences were filtered out because they contained multiple stop codons throughout. The remaining sequences were all pooled and aligned with Muscle (Edgar 2004). Aligned sequences were trimmed with trimal v1.2 using the automated heuristic approach (Capella-Gutiérrez et al. 2009). Unique hotspot sequences were manually selected from the alignments and converted to nucleotide sequences by choosing the most frequent codon in *S. cerevisiae* for each amino acid, based on the CoCoPUTs table for *S. cerevisiae* S288C (TAXID 559292) (Alexaki et al. 2019). For both types of synthetic libraries, the final sequences were generated using custom Python scripts by adding homology regions of 40 bp on either side of the hotspot (S4 Data).

#### Library preparation

Synthetic libraries were ordered as oPools from IDT. Each oPool was resuspended in biomolecular grade water to a final concentration of 0.5 μM for NNK pools and 5 μM for homolog pools, with an additional 10-fold or 100-fold dilution so that all pools were at a final concentration of 0.05 μM for the PCR amplification step. We estimated the total molar amount per tube to be equal to 50 pmol times the number of oligonucleotides (for example for the NNK pool: 9 oligonucleotides x 50 pmol = 450 pmol, resuspended in 900 μL). oPools were amplified with KAPA to add homology regions with sequences directly upstream and downstream of each hotspot, using primers o_022-031. CRISPR-*Cas9* transformations for genome editing were performed as described above, using strains s_003-006 and s_011-014. Transformants were scraped in 5 mL of YPD. The OD_595_ was measured, a glycerol stock was preserved at -70°C and two aliquots of 1 mL each were pelleted for downstream genomic DNA extraction. We estimated a single transformation per condition to yield enough colonies to saturate the expected diversity of variants. Despite a lower number of colonies for FKS1-HS2 (NNK and homolog pools), we managed to get sufficient coverage as we describe in the next section.

#### Pre-screening sequencing runs

For all hotspots except hotspot 3, all libraries (NNK or homologs, obtained from one transformation each) were sequenced to estimate the diversity of variants. For hotspot 3, libraries from four transformations (for each type, NNK and homologs) were sequenced separately. First, genomic DNA was extracted using a standard phenol-chloroform protocol (Amberg et al. 2005). For all hotspots except hotspot 3, PS sites were added by PCR using Q5 and primers o_127-134. Libraries were multiplexed by row-column indexing (Yachie et al. 2016) using KAPA and primers o_135-148. Finally, they were amplified using KAPA and primer pair o_149-150 to add Illumina indexes, SPRI-purified and sequenced in paired-end 300 bp on an Illumina MiSeq system (three runs in total, Institut de Biologie Intégrative et des Systèmes sequencing platform, Université Laval). For hotspot 3, PBS sites were added using Q5 and a touch-down PCR (5 cycles with annealing at 55°C followed by 17 cycles of 2-step amplification, no annealing) with primers o_167-170. For the last step, we added i5/i7 dual barcodes by PCR using KAPA and primers o_187-194/207, SPRI-purified and sequenced in single-end 150 bp on an Aviti system (Institut de Biologie Intégrative et des Systèmes sequencing platform, Université Laval). For all libraries, we confirmed that all expected amino acid variants were present in the initial pools.

#### Inhibition curves

The parental strain BY4741 and strains in which either the hotspot 1 of *FKS1* or the hotspot 1 of *FKS2* was replaced with an hygromycin resistance marker (s_001, s_003 and s_005) were streaked on YPD agar medium supplemented (or not) with hygromycin B. Precultures were prepared from three isolated colonies in 5 mL of YPD medium with appropriate selection and incubated overnight at 30°C with shaking. Precultures were diluted to 1 OD_600_ in sterile water. 3 sterile Greiner 96-well plates were prepared with SC medium supplemented with two-fold dilutions of each of the three echinocandins (for caspofungin, from 0.0156 μg/mL to 0.25 μg/mL; for micafungin, from 0.0625 μg/mL to 1 μg/mL and for anidulafungin, from 0.125 μg/mL to 2 μg/mL) and cultures were inoculated at a final concentration of 0.1 OD_600_. OD_600_ measurements were performed at 30°C every 15 min until a plateau was reached in a Tecan Infinite M Nano (Tecan Life Sciences). The maximum growth rate was transformed into the inhibition coefficient, with an inhibition coefficient of 0 corresponding to the maximum growth rate measured in the absence of drug.

#### Experimental design considerations

Preliminary growth assays using the parental strain BY4741 showed that drug concentrations previously estimated to inhibit growth at 50% or 75% in plates (S12 Fig) were close to completely inhibiting growth in 5 mL tubes, with a doubling time multiplied by more than 7.5 (compared to a culture without drug). Additional tests confirmed that concentrations corresponding to 10% inhibition in plates would be adequate for a screening in 5 mL tubes, with an actual growth inhibition estimated around 82% (corresponding to a doubling time multiplied by roughly 4.5). During our preliminary assays, we noticed that mutating Fks1 or Fks2 could have an effect on cell wall integrity. More specifically, we corroborated previous observations that a functional defect in glucan synthesis could lead to increased levels of chitin in the membrane (Cota et al. 2008). Growth is usually monitored by simple OD_600_ measurements, but in this case, we would risk misestimating cell concentration, which we need to precisely calculate the number of mitotic generations (which is then used to calculate the selection coefficient of each variant). To solve this issue, we measured cell concentration by flow cytometry with a Guava easyCyte HT cytometer (Cytek). Ultimately, we did confirm that concentrations obtained by cytometry do not always correlate with OD_600_ measurements (S4A Fig).

#### Screening

Precultures were prepared in 5 mL of YPD medium supplemented with chloramphenicol to prevent any potential bacterial growth. They were inoculated in triplicates with 100 μL of glycerol stock and incubated overnight at 30°C with shaking. Homologous hotspot variants were spiked in NNK pools by mixing saturated precultures in ratios accounting for the total number of expected variants, based on the measured OD_600_. Later on, read frequencies confirmed that the pools were mixed in the appropriate proportions. The mixed precultures were diluted to 1.5 OD_600_ in sterile water and used to inoculate 5 mL tubes containing SC medium supplemented with either sterile water, 0.022 μg/mL caspofungin, 0.15 μg/mL micafungin or 0.22 μg/mL anidulafungin, at a final concentration of 0.15 OD_600_. Additionally, pellets of 1 mL of mixed precultures were prepared and stored at -70°C until genomic DNA extraction (“t0/input” samples). Cultures were incubated at 30°C with shaking until they reached approximately 1 OD_600_, at which point they were adjusted to 1.5 OD_600_. Pellets of 1 mL were prepared and stored at -70°C (“t1 samples”) and fresh cultures were inoculated at a final concentration of 0.15 OD_600_ for a second round of selection. After diluting the t1 cultures at 0.6 OD_600_, cell concentrations were measured by flow cytometry. After reaching approximately 1 OD_600_, t2 cultures were processed similarly to obtain “t2/output” samples (except for hotspot 3 cultures supplemented with drugs, which were incubated overnight). The number of mitotic generations per selection round for each sample was calculated from cytometry measurements of cell concentrations by subtracting the logarithm of values before/after selection and dividing by the logarithm of 2. The total number of generations over both rounds of selection was obtained by summing up the numbers obtained for each round.

#### Genomic DNA extraction

Genomic DNA was extracted from t0 and t2 samples using the DNeasy Blood and Tissue kit (Qiagen), with the regular format for a couple of samples and all hotspot 3 samples, and with the DNeasy 96 protocol for the remaining samples. We followed the kit’s protocol except for the following modifications: initial lysis was performed with 6 units of zymolyase for 1h at 37°C, spheroplasts lysis was performed for 30 min at 56°C, RNase treatment was performed immediately after elution by adding 4 μL of 10 mg/mL RNase A and incubating 1h at 37°C. Samples were then SPRI-purified with a bead ratio of 0.4 and ultimately eluted in 50 μL of 10 mM Tris-HCl.

#### Post-screening sequencing runs

Two preliminary post-screening sequencing runs were performed from 10 and 4 samples, respectively. The first run was performed on an Illumina MiSeq system, whereas the second one was performed on an Illumina iSeq system. For the MiSeq run, PS sites were added using homemade Phusion polymerase and a touch-down PCR (5 cycles with annealing at 50°C followed by 15 cycles with annealing at 68°C), with the same primers as before (o_127-134).

The rest of the protocol was similar to what was done previously (row-column indexing and final amplification using KAPA, SPRI purification and sequencing in paired-end 300 bp at the Institut de Biologie Intégrative et des Systèmes sequencing platform). For the iSeq preliminary run and both ultimate runs (NovaSeq for hotspot 1 and 2, Aviti for hotspot 3), PBS sites were added similarly to what was done with the pre-screening Aviti sequencing run of hotspot 3, i.e. using Q5 and a touch-down PCR (5 cycles with annealing at 55°C followed by 17 cycles of 2-step amplification, no annealing) with primers o_151-170. For the last step, we added i5/i7 dual barcodes by PCR using KAPA and primers o_171-237. iSeq sequencing was performed in paired-end 150 bp at the CHUL sequencing platform, Université Laval. For both ultimate runs, all libraries were quantified in four technical replicates using the AccuClear Ultra High Sensitivity dsDNA Quantitation kit (Biotium). Quantified libraries for the NovaSeq run were manually pooled in four subpools of different concentrations (1X, 4X, 50X, 200X) depending on the concentration of individual libraries to optimize the pipetting volume. Overall, we aimed at an output of 800 million reads, with a coverage of 237X for every library. The four subpools were finally assembled into a single pool at a final concentration of 14 nM. Libraries were sequenced in paired-end 150 bp on an Illumina NovaSeq 6000 S4 system at the Centre d’expertise et de services Génome Québec. Hotspot 3 libraries were sequenced in single-end 150 bp on an Aviti system (Institut de Biologie Intégrative et des Systèmes sequencing platform, Université Laval).

### Computational analyses of DMS data

The following sections describe the main steps of each computational analysis. Technical details and all scripts are available online on the dedicated GitHub repository: https://github.com/Landrylab/Durand_et_al_2026. Jobs were run on the IBIS servers.

#### Quality control

Initial assessment of sequencing data showed that the number of obtained reads was almost consistently larger than the number of expected reads (3 times on average), with very little variability between samples. Most reads could be confidently attributed to an expected variant, resulting in an overall coverage of more than 600X across all conditions.

#### Processing sequencing data

The forward reads of DMS sequencing data were analyzed with gyōza v1.2.1 (Durand et al. 2026). Data from the MiSeq pre-sequencing runs were demultiplexed using a custom script with several consecutive rounds of demultiplexing: a first round with DemuxFastqs (now fqtk) from Fulcrum Genomics to demultiplex forward plate barcodes by specifying the expected read structure (5M9B+T 5M+T), a second round also with DemuxFastqs to demultiplex reverse plate barcodes (+T 9B+T) and a third round with cutadapt to demultiplex row-column indexes. All other data, including from the main sequencing runs, were already demultiplexed and could be processed directly by gyōza. Details of the configuration are available on the dedicated GitHub repository.

Only variants sequenced more than 10 times in all replicates were labeled “high confidence” and used in downstream analyses. gyōza uses a standard log-ratio method to calculate selection coefficients for each amino acid sequence. This includes a normalization with the number of mitotic generations (g; previously obtained from cytometry measurements of cell concentrations before and after each round of selection), as well as a normalization with the median log2 fold-change of silent mutants, except for the wild-type nucleotide sequence. The reason why the wild-type nucleotide sequence is omitted from this normalization is because it is overabundant in our oPool design. The Spearman’s rank correlation coefficient was calculated for pairwise comparisons of replicates (S6 Fig).

#### Classification of mutational effects

The multimodal distribution of selection coefficients for each condition was used to classify variants, using a Gaussian mixture model (implemented in Python with the scikit-learn package). For each combination of mutated hotspot and compound (drug or control condition), we first optimized the model by calculating the Akaike Information Criterion (AIC) and the Bayes Information Criterion (BIC) for up to five components (S7 Fig). We then ran the model using the number of components that minimized the BIC: 3. Based on the mean of each distribution, we converted the labels predicted by the model to biologically relevant effects on fitness (either deleterious, slightly deleterious or WT-like in the absence of drug i.e. effect on protein function, or WT-like, intermediate or resistant in the presence of drug i.e. effect on resistance). To resolve overlaps between distributions, we set thresholds 2.5 standard deviations above and below the mean of the narrowest distributions (more details in S7 Fig). Finally, we collapsed bins to simplify the classification: variants were considered either sensitive or resistant (non-sensitive) to a drug and either deleterious or non-deleterious in the absence of drug.

### Validations of mutational effects

Two validation assays were performed similarly: the first one with 28 single mutants of Fks1-HS1 to validate the DMS and the second one to validate the docking, using mutants outside the hotspots (and D691V in hotspot 3), as well as F639S, F639D and R645D in hotspot 1 from the first assay to be used as controls. We also included the parental strain BY4741 and strains in which the wild-type sequence was reinserted in the appropriate deletion background. Strains were grown overnight in 1 mL of YPD in a 96-deep-well plate at 30°C with shaking. Precultures were diluted to 1 OD_600_ in sterile water and inoculated at a final concentration of 0.1 OD_600_ in sterile Greiner 96-well plates with SC medium supplemented with either sterile water or echinocandin. For the assay designed to validate the DMS assay, we used 0.058 μg/mL caspofungin or 0.44 μg/mL micafungin or 0.63 μg/mL anidulafungin. The concentrations were chosen because they correspond to 80% inhibition based on our inhibition curves (S12 Fig), which is approximately what we estimated was the actual inhibition coefficient for the DMS screening (82 %). For the assay designed to validate the docking, we used concentrations that completely inhibit WT growth to ensure discrimination of resistant vs sensitive mutants: 0.232 μg/mL caspofungin or 1.76 μg/mL micafungin or 2.52 μg/mL anidulafungin. OD_600_ measurements were performed at 30°C every 15 min until a plateau was reached in a BioTek Epoch 2 Microplate Spectrophotometer (Agilent). For each mutant, we calculated the area under the curve (AUC) using the composite trapezoidal rule (implemented with the trapezoid function from the Python package numpy) and converted it to a log2 fold-change, normalizing with the value obtained with the parental strain. The recreated wild-type control had a fitness effect similar to that of the parental strain, confirming that the observed fitness effect is largely a consequence of the introduced mutation, rather than that of one or several putative off-target mutations.

Correlation between the obtained relative fitness values and the selection coefficients from the DMS assay was evaluated by calculating the Spearman’s rank correlation coefficient (S8 Fig). The level of micafungin resistance of only two mutants (V641W and L642K) was underestimated in the DMS experiment. For both, a corrected DMS score was obtained by linear regression, which allowed us to re-classify the variants appropriately.

### Molecular modeling

#### Fks1 structure preparation

The 3.4 Å cryo-EM structure of the Fks1 protein (PDB ID: 7XE4 (Hu et al. 2023)), was used. The ten missing loop segments (residues 244-278, 475-487, 799-805, 897-931, 1159-1167, 1247-1266, 1419-1435, 1516-1554, 1627-1637, and 1698-1723) were reconstructed using the GalaxyFill tool (Coutsias et al. 2004) within the CHARMM-GUI framework (Jo et al. 2008). The reconstructed Fks1 was validated using the Molecular Operating Environment’s (MOE) structure preparation tool (2022. 02 Chemical Computing Group ULC 2024), using the CHARMM27 force-field. The protonation states of residues were determined at pH 7.0, employing the APBS and PDB2PQR tools (Jurrus et al. 2018) under default CHARMM force-field parameters.

#### Fks1-membrane system construction

The position of the resulting Fks1 in the membrane was predicted using the PPM 3.0 (N-ter: in; fungi plasma membrane) server (Lomize et al. 2022). The Membrane Builder tool from the CHARMM-GUI server (Wu et al. 2014) was used to construct the protein-membrane system. This preparation incorporated post-translational modifications as specified by the cryo-EM structure article (Hu et al. 2023), including termini truncation with acetylation (ACE) and amidation (CT3) at the N- and C-termini, respectively, the disulfide bonds (658-669, 1328-1345), and the addition of a di-N-acetylglucosamine moiety (β-D-GlcNAc (1→4) β-D-GlcNAc) at N1849. A lipid bilayer mimicking the asymmetrical fungal plasma membrane’s composition was constructed (Pogozheva et al. 2022), consisting in the following lipid compositions : DYPC, YOPC, POPE, PYPE, YOPE, POPI, POPS, YOPA, ERG, and PI-Ceramide (18:1,16) with (->2) linked D-mannose, and the ratio of 8:3:3:1:3:5:7:3:49:54 for the outer leaflet, and 10:6:10:5:6:20:31:8:4:0 for the inner leaflet, respectively, for a total of 907 lipids. The system was neutralized and the ionic concentration was 0.15 mM NaCl. The final systems measured approximately 173 x 173 x 182 Å on the XYZ dimensions, and water molecules were added to create a 30 Å layer on either side of the membranes along the Z axis, resulting in systems of 507,096 atoms.

#### Simulation details and parameters

Three independent 1000-ns Molecular Dynamics (MD) trajectories were conducted to allow protein relaxation. These simulations were performed with the software NAMD (Phillips et al. 2020), using the CHARMM36m force field parameters (Huang et al. 2017) and TIP3P waters (Durell et al. 1994). Simulations were carried out at 303.15 K under isothermal-isobaric (NPT) ensemble conditions with a 2-fs time step and periodic boundary conditions. Langevin damping with a coefficient of 1 ps^−1^ was used to maintain constant temperature, while pressure was controlled by a Nosé-Hoover Langevin piston at 1 atm. The length of the bonds between hydrogen and heavy atoms was constrained using SETTLE (Miyamoto and Kollman 1992) for water molecules and SHAKE (Ryckaert et al. 1977) for all other molecules. Cut-off distance for the short-range electrostatics and the Lennard-Jones interactions was 12 Å and a switching function was used from 10 to 12 Å. Nonbonded pair lists were updated every 10 steps. The particle mesh Ewald (PME) method (Darden et al. 1993) was applied for the calculation of long-range electrostatic interactions at each integration step using a sixth-order interpolation and a grid spacing of ∼1 Å. The non-bonded pair lists were updated every ten steps. The CHARMM-GUI Membrane Builder protocol for system equilibration was applied (Wu et al. 2014). The last frame of each trajectory is shown in S1 Fig.

### Sequence and structure comparisons of Fks homologs

Sequence comparisons were conducted in MEGA X (Kumar et al. 2018) and inferred by using the maximum likelihood method based on the LG matrix-based model (Le and Gascuel 2008). Fks homologous sequences were aligned and trimmed as described above. Initial tree(s) for the heuristic search were obtained automatically by applying Neighbor-Join and BioNJ algorithms to a matrix of pairwise distances estimated using the JTT model, and then selecting the topology with the superior log likelihood value. A discrete Gamma distribution was used to model evolutionary rate differences among sites (5 categories (+*G*, parameter = 0.7192)). The rate variation model allowed for some sites to be evolutionarily invariable ([+*I*], 2.57% sites).

The Fks1 structure from *C. albicans* was retrieved from the AlphaFold database (AFDB accession AF-A0A1D8PCT0-F1) (Varadi et al. 2024). Fks1 structures from *N. glabratus*, and *C. auris* were predicted using AlphaFold v2.3.2 (Jumper et al. 2021). Structural alignment was obtained with US-align (Zhang et al. 2022). Conservation scores computed with Jalview were used to visualize conservation across the aligned structures (Livingstone and Barton 1993), using ChimeraX (Pettersen et al. 2021).

### Softwares, packages and versions

The following Python (v3.13) packages were used: snakemake v9.14.0 (Mölder et al. 2021) to run gyōza v1.2.1 (Durand et al. 2026), pandas v2.3.3 (McKinney 2010), matplotlib v3.10.7 (Hunter 2007), numpy v2.3.5 (Harris et al. 2020), papermill v2.6.0, scikit-learn v1.7.2 (Pedregosa et al. 2011), scipy v1.16.3 (Virtanen et al. 2020), seaborn v0.13.2 (Waskom 2021), logomaker v0.8.7 (Tareen and Kinney 2020) and a forked version of upsetplot. Molecular models were manipulated in PyMOL v2.5.5 (Schrödinger, LLC 2015) or ChimeraX v1.8 (Pettersen et al. 2021).

## Acknowledgments

We thank Anna Fijarczyk, François D. Rouleau and Stéphane Larose for bioinformatic assistance, Jordan Jalbert-Ross with technical assistance on preliminary assays, Philippe C. Després, François D. Rouleau and Soham Dibyachintan for critical reading of the manuscript and the whole team for feedback and emotional support on a regular basis throughout the project.

## Funding

Canadian Institutes of Health Research (CIHR) (Foundation grant 387697, CRL)

Genome Canada and Genome Quebec (grant 6569, CRL)

Canada Research Chair in Cellular Systems and Synthetic Biology (CRL)

Natural Sciences and Engineering Research Council of Canada (NSERC) (Grant RGPIN-2022-04721, PL)

Fonds de Recherche du Québec - Santé (FRQS) postdoctoral fellowship (RD) NSERC postdoctoral fellowship (RD)

## Author contributions

Conceptualization: RD, AGT, PL, CRL

Methodology: RD, AGT, MG, AP, AKD, PL, CRL

Software: RD, AGT, MG, AP, PL

Validation: RD, MG, AP, AKD, PL

Formal analysis: RD, AGT, MG, AP, PL

Investigation: RD, AGT, MG, AP, AKD, PL

Resources: PL, CRL

Data curation: RD, MG, AP

Writing - Original draft: RD, AGT, MG

Writing - Review and editing: RD, AGT, PL, CRL

Visualization: RD, AGT, MG, AP, PL

Supervision: RD, AGT, AP, AKD, PL, CRL

Project administration: PL, CRL

Funding acquisition: PL, CRL

## Data and materials availability

All data are available in the main text or the supplementary materials. Sequencing data are available at the NCBI Sequence Read Archive (SRA) under BioProject PRJNA1126009: https://www.ncbi.nlm.nih.gov/sra/PRJNA1126009. All data and code to reproduce results and figures for the DMS and the machine learning sections are available at https://github.com/Landrylab/Durand_et_al_2026. All strains listed in S4 Data are available and can be requested by email at.

## Supplementary Materials

S1 to S12 Figures

S1 to S4 Data

