## Supplemental Material for "Mutational landscape and molecular bases of echinocandin resistance in *Saccharomyces cerevisiae*"

**The PDF file includes:**

S1 to S12 Figures

**Other Supplementary Materials for this manuscript include the following:**

S1 to S4 Data

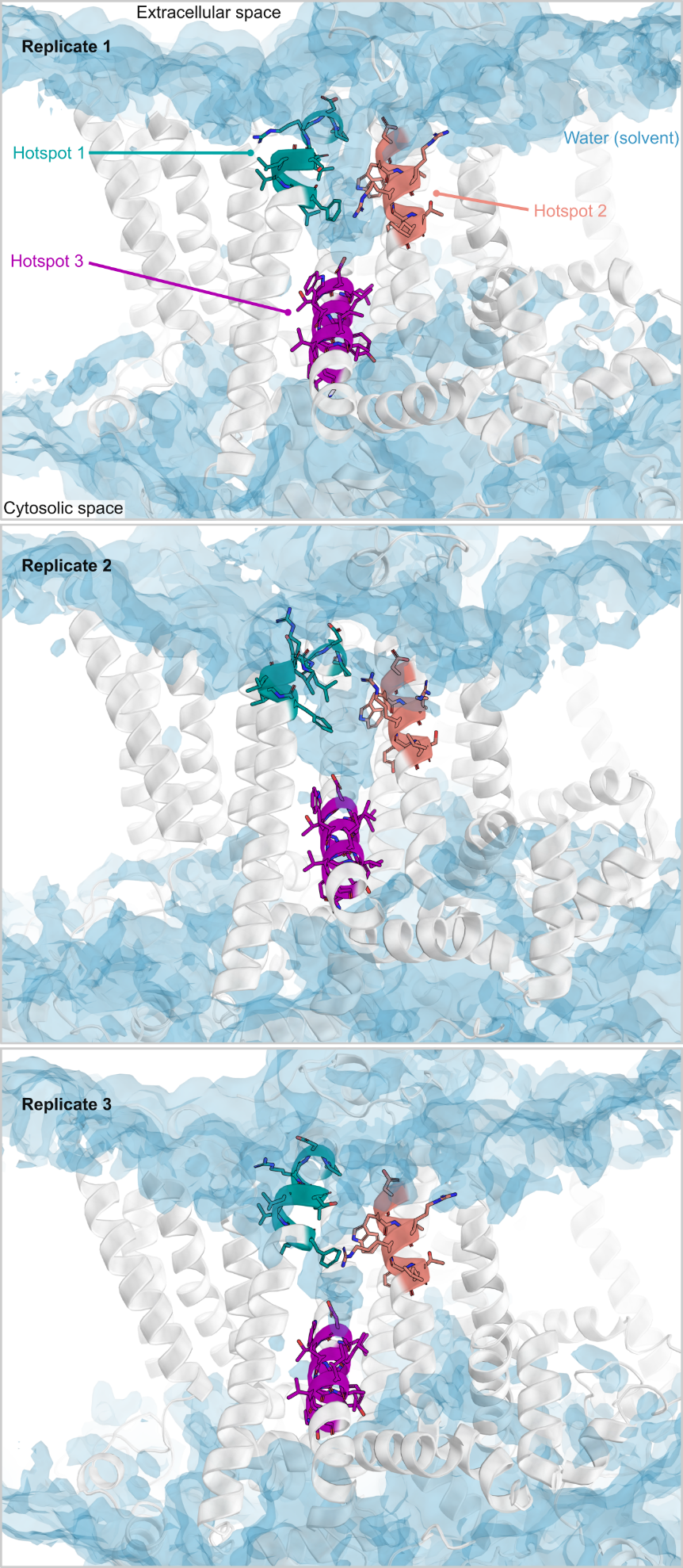

**S1 Figure. Membrane-embedded water pockets connecting Fks1 hotspots from three independent trajectories.** Water occupancy maps (blue surface, isocontour 0.2), each computed from one of three independently constructed MD trajectories of Fks1 (white ribbon) in the absence of echinocandin revealed a water-filled pocket in the transmembrane region. The average of these three replicates is shown in Fig. 1. This feature was located primarily between hotspot 1 (dark cyan) and hotspot 2 (salmon), extending toward hotspot 3 (purple). Figures were generated using PyMOL v2.5.5 (Schrödinger, LLC 2015).

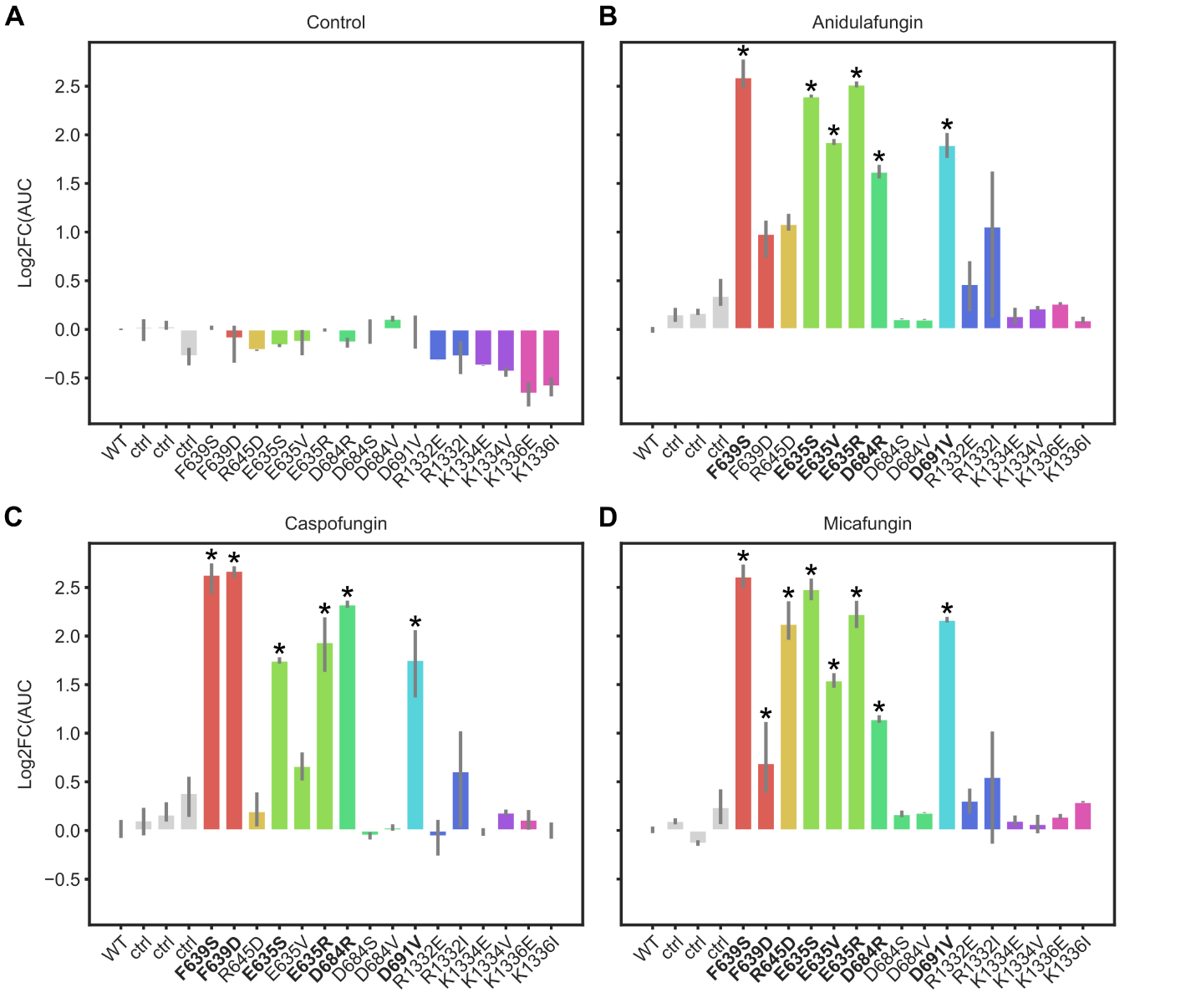

**S2 Figure. Individual growth measurements for mutants outside the hotspots.**

13 single mutants of Fks1 were recreated in the parental strain and used in a microplate growth assay using drug concentrations completely inhibiting WT growth. Controls include the WT (parental strain), strains obtained by reinserting the WT hotspot in the same deletion background as the other mutants (labeled “ctrl”) and strains corresponding to 3 Fks1-HS1 mutants used in the validation assay featured in S8 Fig. Relative growth corresponds to the log2-transformed area under the curve (Log2FC(AUC), calculated on 25h) from 2-3 replicate colonies, normalized by the parental strain. Error bars indicate the full range of values. The baseline for resistance appears to vary from one drug to another, as suggested by the growth measured for F639D, which is specifically sensitive to anidulafungin. Accordingly, R645D, which is specifically resistant to micafungin, is an additional indication that mutants tested in this assay can be considered resistant to anidulafungin if they grew more than F639D or R645D. On the other hand, a mutant growing as little as F639D in micafungin is most likely resistant. To simplify interpretations, we labeled the bars of resistant mutants with “*” and highlighted their labels in bold.

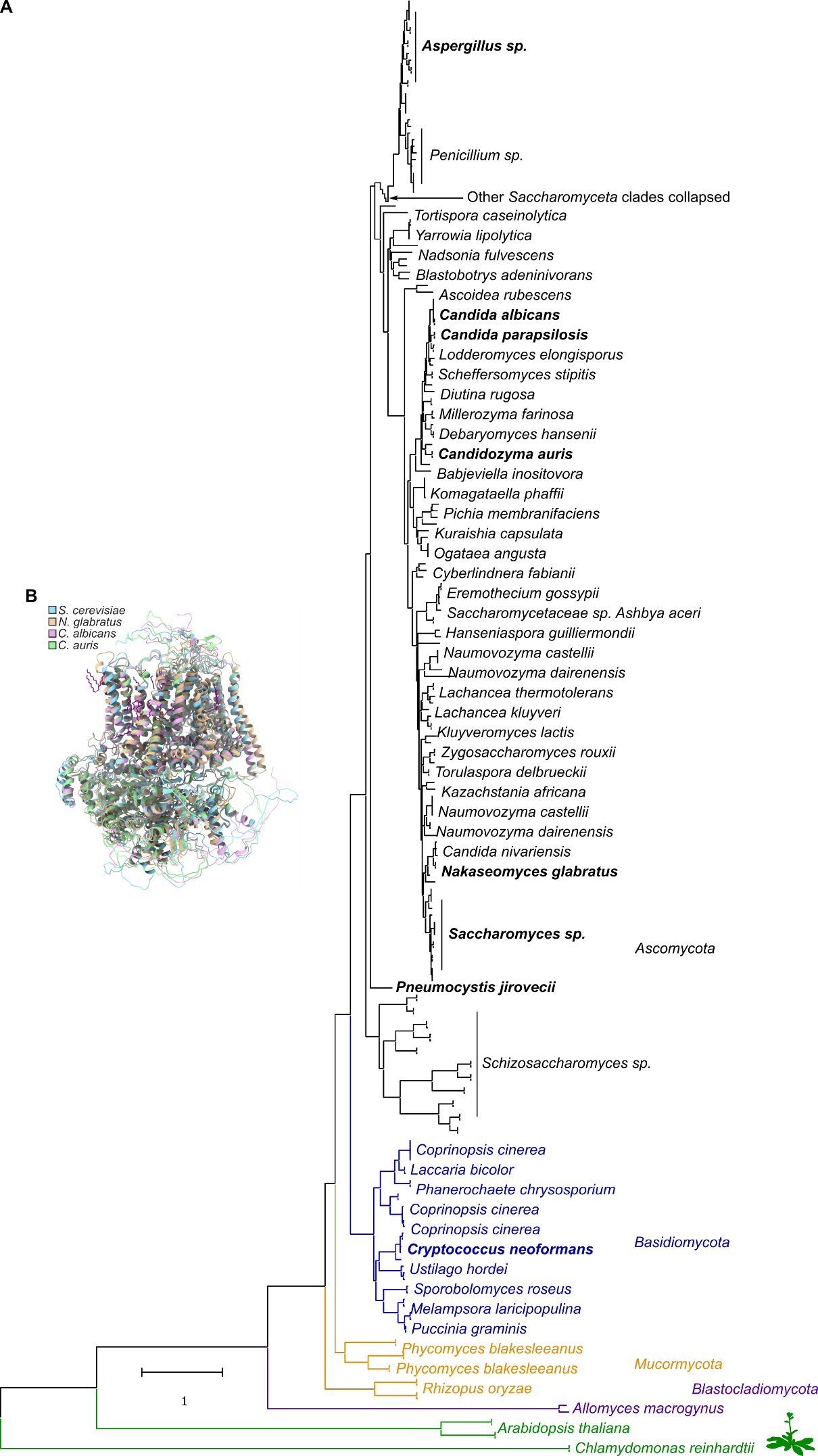
 **S3 Figure. Conservation of Fks homologs.**

(**A**) Maximum likelihood phylogenetic analysis of all Fks homologs. Tree is drawn to scale, with branch lengths measured in the number of substitutions per site over 1,731 amino acid positions (410 amino acid sequences). Homologs found in the *Plantae* kingdom (green) were used to root the tree (triple pun intended). *Ascomycota*, *Basidiomycota* and *Mucormycota* are colored in black, blue and orange, respectively. Several key species are highlighted in bold (*S. cerevisiae* and clinically relevant pathogens *C. auris*, *N. glabratus*, *C. parapsilosis*, *C. albicans*, *A. fumigatus*, *C. neoformans* and *P. jirovecii*). (**B**) Alignment of the cryo-EM structure of Fks1 from *S. cerevisiae* (PDB: 7XE4 (Hu et al. 2023)) and the AlphaFold predicted structures (see methods) of Fks1 from *C. albicans* (AF-A0A1D8PCT0-F1), *N. glabratus* and *C. auris*, visualized using UCSF ChimeraX v1.8 (Pettersen et al. 2021).

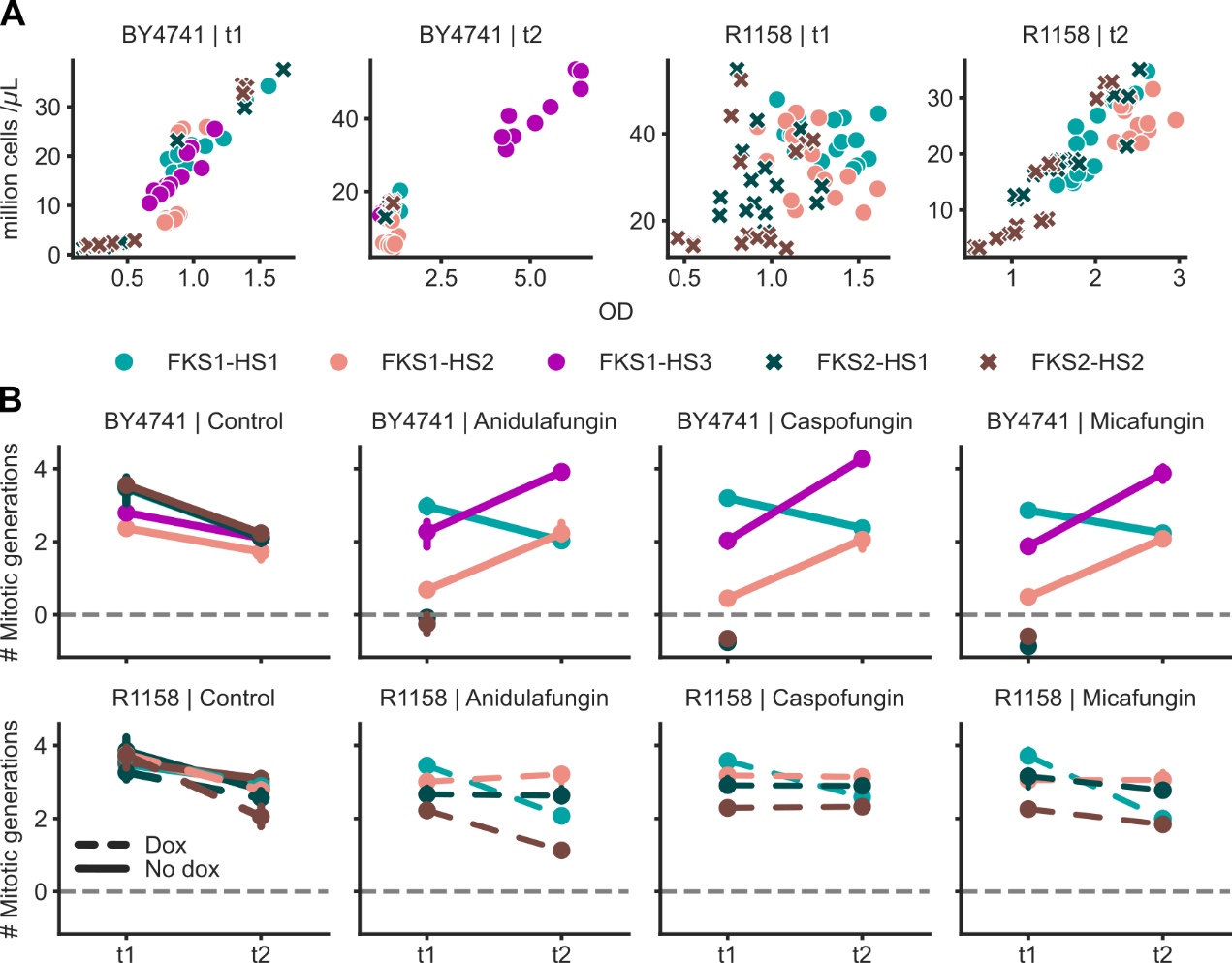

**S4 Figure. Growth measurements during the DMS screen.**

(**A**) Relationship between cell concentrations measured by cytometry and optical density (OD_600_) measurements at t1 and t2 for both backgrounds (BY4741, native expression of paralog; R1158, paralog repressed). (**B**) Number of mitotic generations calculated from flow cytometry measurements. For each sample, we subtracted the logarithms of cell concentrations at t1/t0 (“t1”) or t2/t1 (“t2”) and divided by the logarithm of 2. The sum of both values (total number of generations over both rounds of selection) was used to normalize fold-changes of allele frequencies to obtain selection coefficients. For all conditions involving repression of the paralog, doxycycline (“Dox”) was added to the medium (except for the “No dox” control).

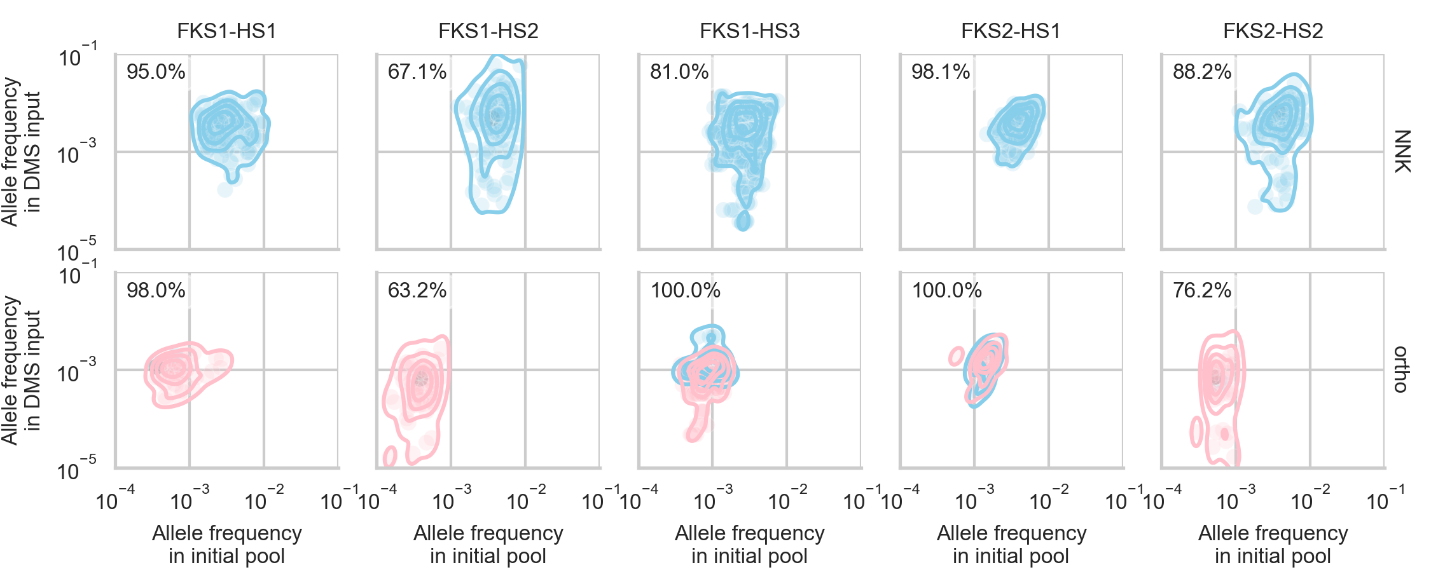

**S5 Figure. Comparison of allele frequencies before/after overnight culture.**

Density plots indicating the allele frequencies of each amino acid variant in the DMS input and in the initial pools of NNK and homologous hotspot variants, with native paralog expression (BY4741 background). 5 contour levels indicate iso-proportions of the density for variants that differ from the wild-type by a single amino acid change (in blue) or more (pink). Percentages indicate the proportion of unique variants sequenced more than 10 times in both the initial pools and the DMS input, relative to the total number of unique expected variants.

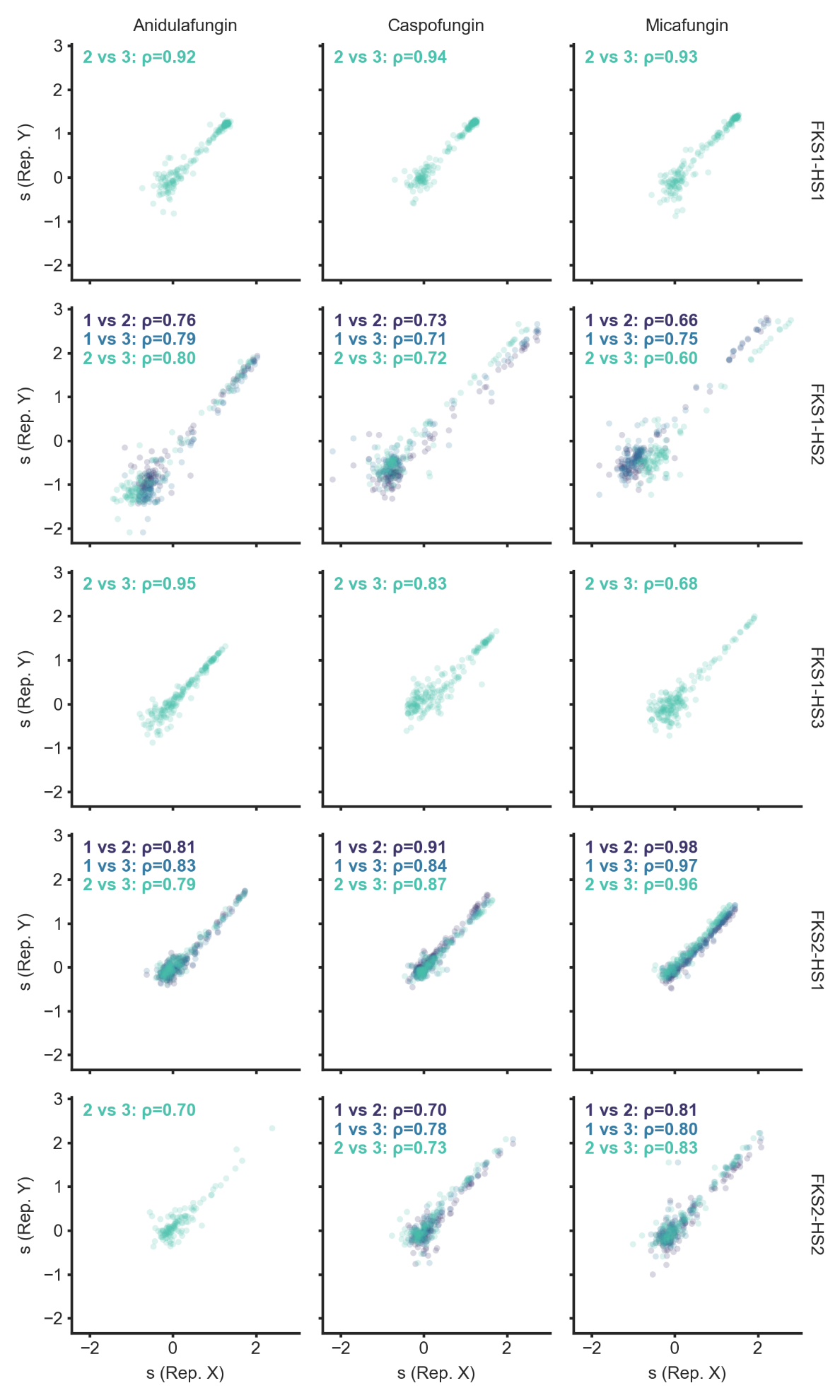

**S6 Figure. Correlation between selection coefficients from replicates.**

Pairwise comparisons of selection coefficients (s) between replicates (Rep.) for *FKS1* hotspots with native expression of *FKS2* (BY4741 background) or *FKS2* hotspots with repressible expression of *FKS1* (R1158 background). Replicates correspond to equivalent libraries cultured separately overnight (i.e. pre-cultures in separate 5 mL culture tubes), with each pre-culture used to inoculate every culture corresponding to different drug conditions. Each data point corresponds to a single amino acid sequence. In all downstream analyses, the median across replicates was used. Empty panels indicate one replicate was lost during gDNA extraction. For each pairwise comparison, the Spearman’s rank correlation coefficient is indicated.

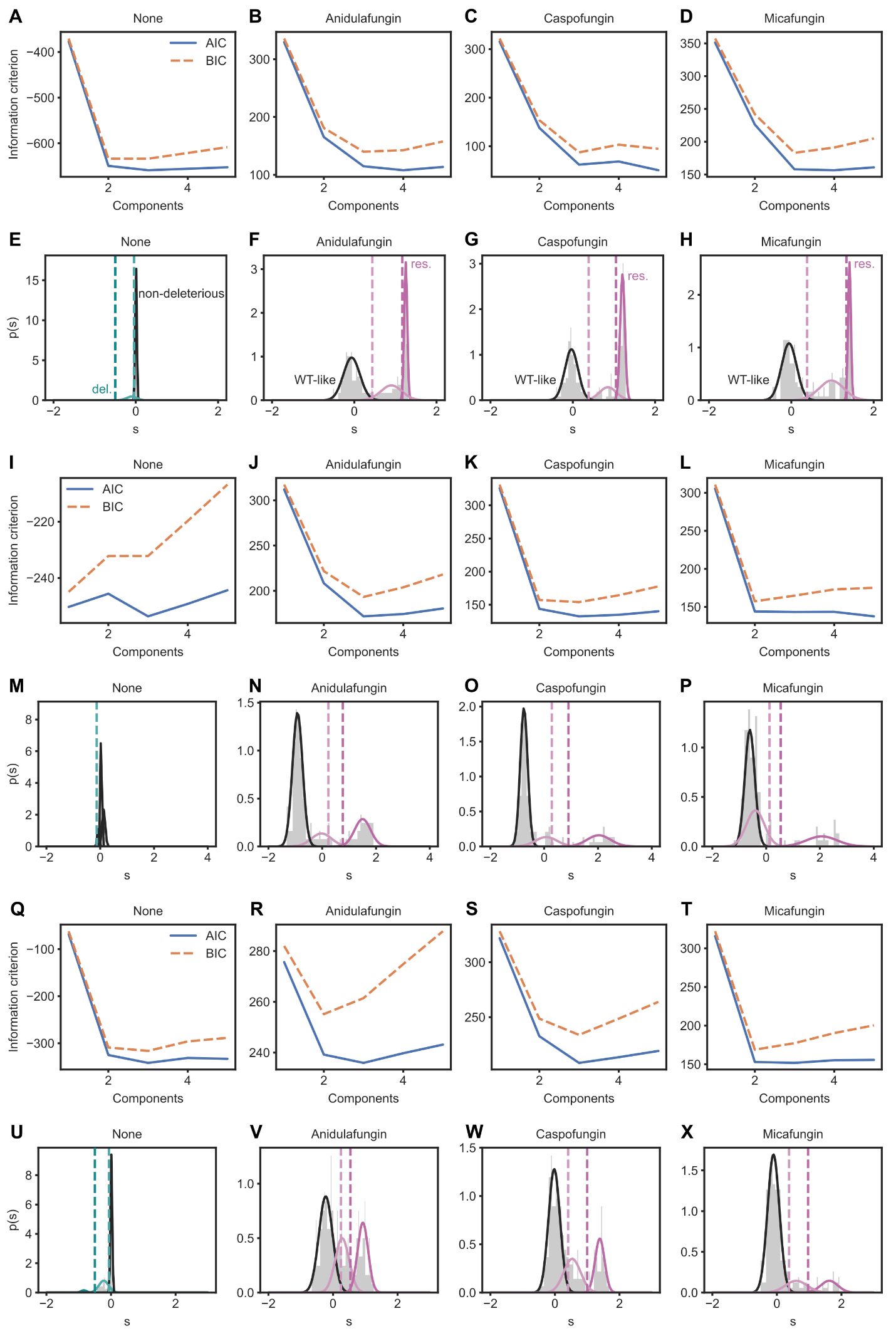

**S7 Figure. Classification of mutational effects based on the distribution of selection coefficients.**

Classification of FKS1-HS1 (**A-H**), FKS1-HS2 (**I-P**) and FKS1-HS3 (**Q-X**) mutants using a Gaussian mixture model. Panels A-D, I-L and Q-T show the optimization step, which estimates the number of components by minimizing the Bayes Information Criterion (BIC) value. As a result, the model was run with 3 components. Panels E-H, M-P and U-X show the densities for each component/mode, as predicted by the model. The density of the full (multimodal) distribution of selection coefficients for each condition is indicated in gray. Based on the mean of each (colored) predicted distribution, the three components were converted to three phenotypic effects: either deleterious (lowest mean), slightly deleterious or non-deleterious in the absence of drug (impact on protein function), or WT-like, intermediate phenotype or resistant (highest mean) in the presence of drug (impact on resistance). To resolve overlaps, we set thresholds 2.5 standard deviations above or below the narrowest distributions (colored dashed lines in panels E-H, M-P and U-X). FKS1-HS1 variants with a selection coefficient between the WT-like mean + 2.5 standard deviations and the resistant mean - 2.5 standard deviations were classified as intermediate. For FKS1-HS2, because the narrowest distribution was the intermediate, we set the upper threshold at intermediate mean + 2.5 standard deviations instead. In the control condition, variants with a selection coefficient between -0.5 and the WT-like mean - 2.5 standard deviations were classified as slightly deleterious. Ultimately, the classification was collapsed into a binary one: either deleterious or non-deleterious (initially slightly deleterious or non-deleterious) in the absence of drug, and either sensitive or resistant (non WT-like).

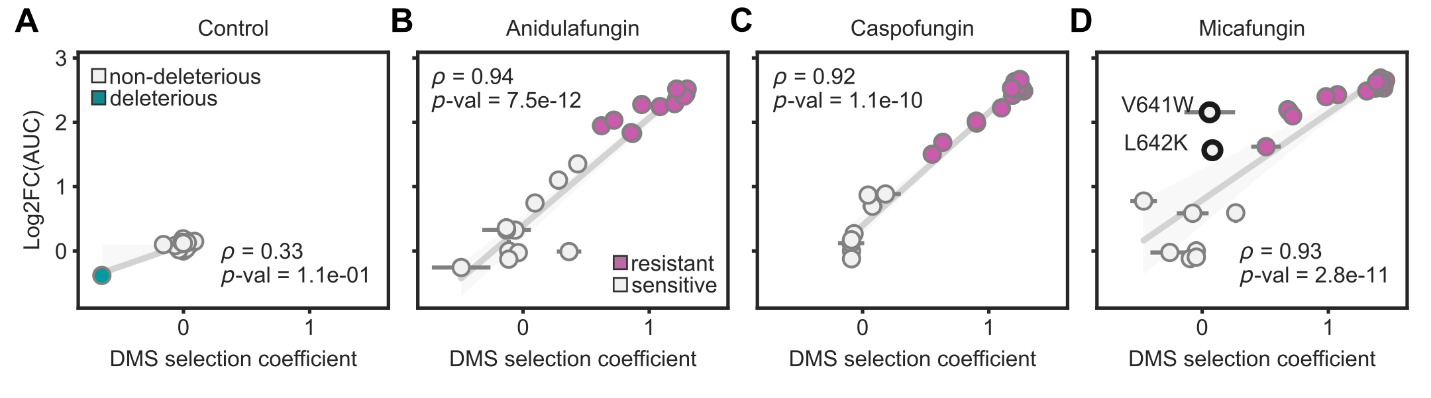
**S8 Figure. Individual growth measurements for Fks1-HS1 mutants and comparison with effects from the DMS experiment.**

28 single mutants of FKS1-HS1 were recreated in the parental strain and used in a microplate growth assay using drug concentrations inhibiting 80% of WT growth. For all four conditions (control, **A**; anidulafungin, **B**; caspofungin, **C**; micafungin, **D**), two additional data points correspond to the WT: the parental strain and the control for which the WT hotspot was reinserted similarly to the other mutants. Relative growth corresponds to the log2-transformed area under the curve (Log2FC(AUC), calculated on 40h) from up to two replicate colonies, normalized by the parental strain. Error bars indicate the full range or first and third quartiles for relative growth and DMS selection coefficient, respectively (across replicates in both cases). For each condition, correlation between the two scores was estimated by calculating the Spearman’s rank correlation coefficient (ρ) and the corresponding *p*-value. The slope and intercept of the linear regression (statsmodels ordinary least squares) were used to calculate an “inferred” DMS selection coefficient. For the two mutants highlighted in panel D, this score was used rather than the initially underestimated selection coefficient to appropriately re-classify the mutants in this specific condition. The classification is reflected in the color code (except for the highlighted mutants in panel D which were re-classified as resistant).

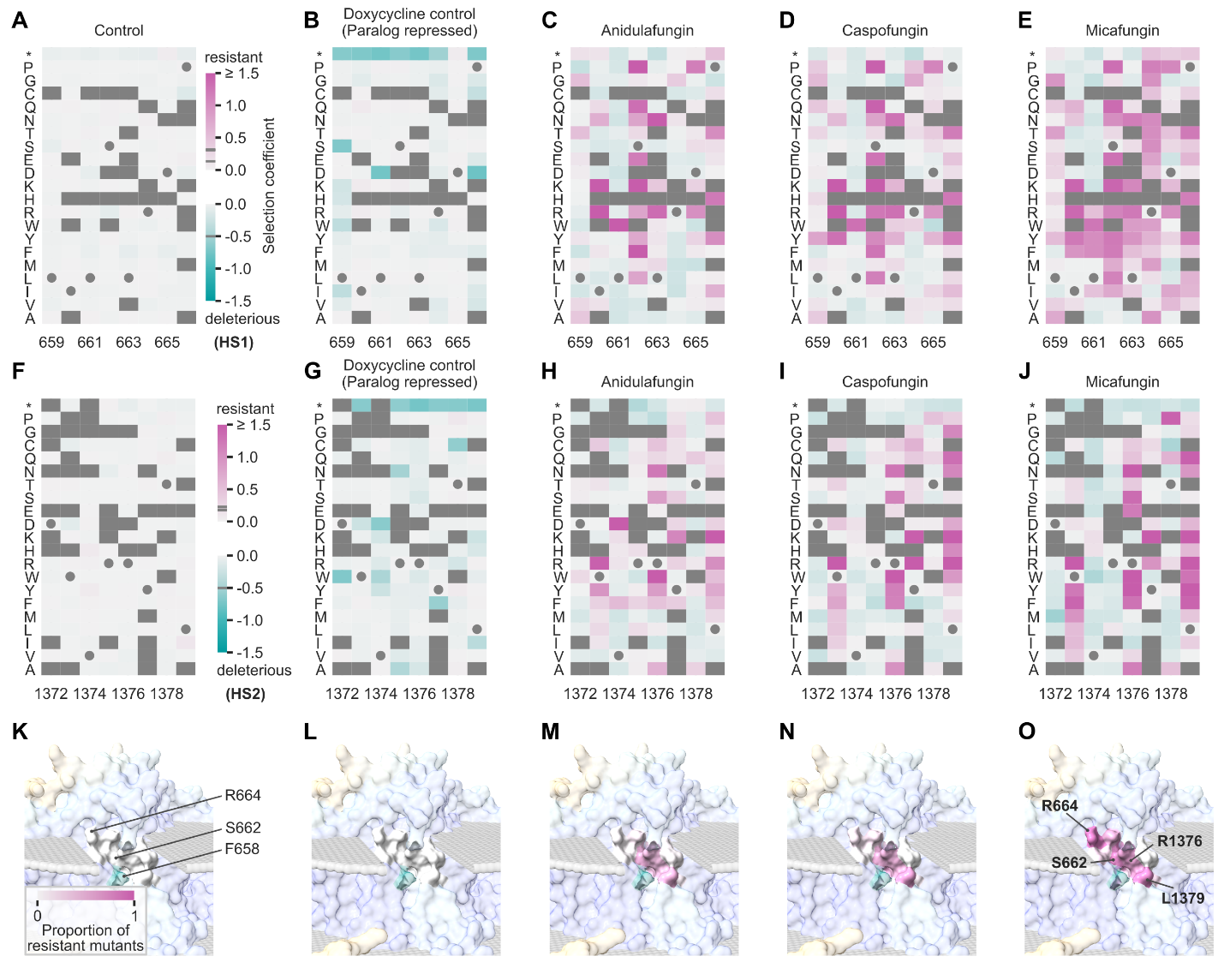

**S9 Figure. Mutational effects in Fks2.**

(**A-J**) Heatmaps showing the selection coefficient (median of synonymous variants) for each single mutant of hotspot 1 (**A-E**) and hotspot 2 (**F-J**) of Fks2, across conditions: control (**A, F**), doxycycline alone (**B, G**), anidulafungin + doxycycline (**C, H**), caspofungin + doxycycline (**D, I**) and micafungin + doxycycline (**E, J**). Doxycycline induces the repression of the paralog *FKS1*. The wild-type sequence is indicated by gray circles. Gray squares indicate mutants that were not sequenced with sufficient coverage before screening. (**K-O**) For each condition (same order as previous panels), both hotspots in the AlphaFold2-predicted structure of Fks2 are colored according to the proportion of mutants classified as resistant at each position (relative to the number of mutants for which we were able to obtain a selection coefficient), with key residues highlighted. Semi-opaque shades of blue and yellow used to color the structure surface indicate high and low levels of confidence in the structure prediction, respectively.

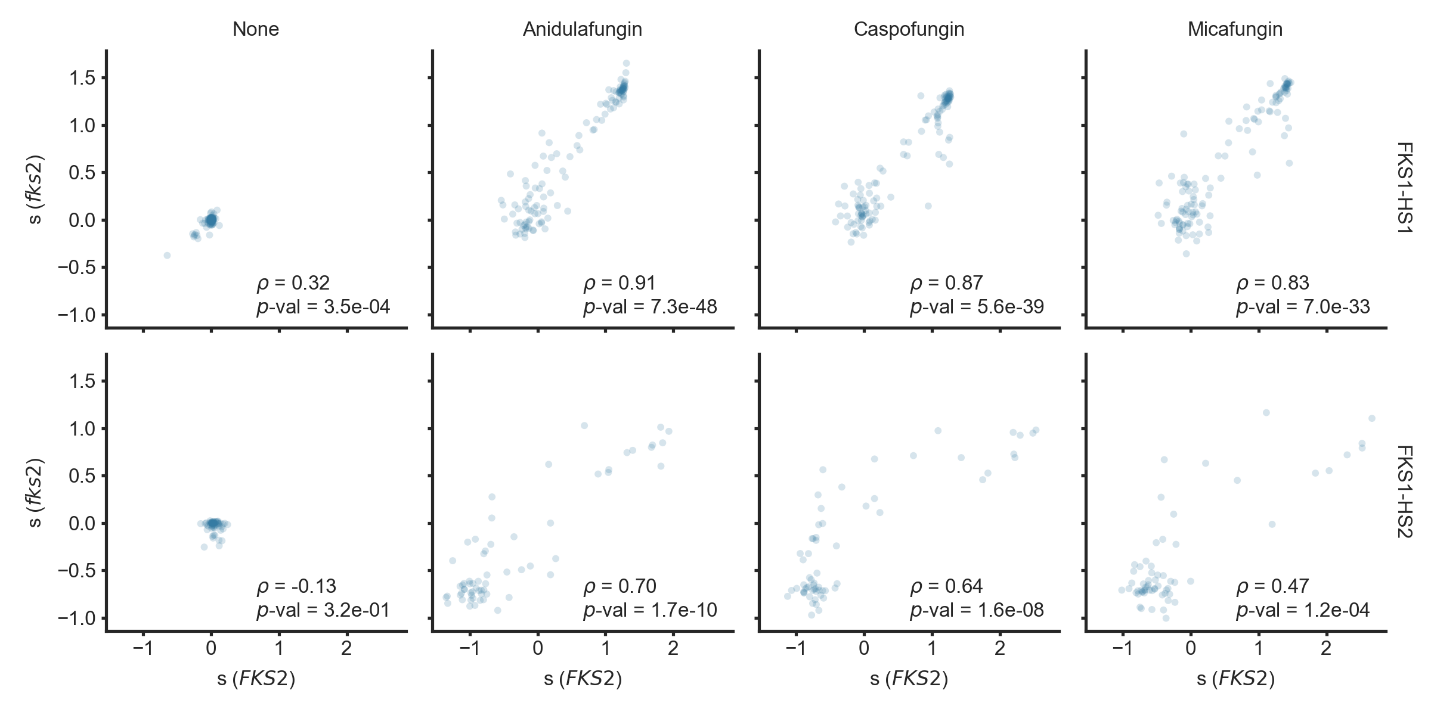

**S10 Figure. Mutational effects with or without *FKS2* repression.**

Fks1 libraries were built in two different backgrounds: BY4741 and R1158, to assess mutational effects in hotspot 1 (top row) and hotspot 2 (bottom row) with (*FKS2*) or without (*fks2*) native expression of the paralog, respectively. In the latter case, we used doxycycline (in addition to the indicated echinocandin). Each data point corresponds to a single amino acid sequence. Spearman’s rank correlation coefficients (ρ) and *p*-values are indicated.

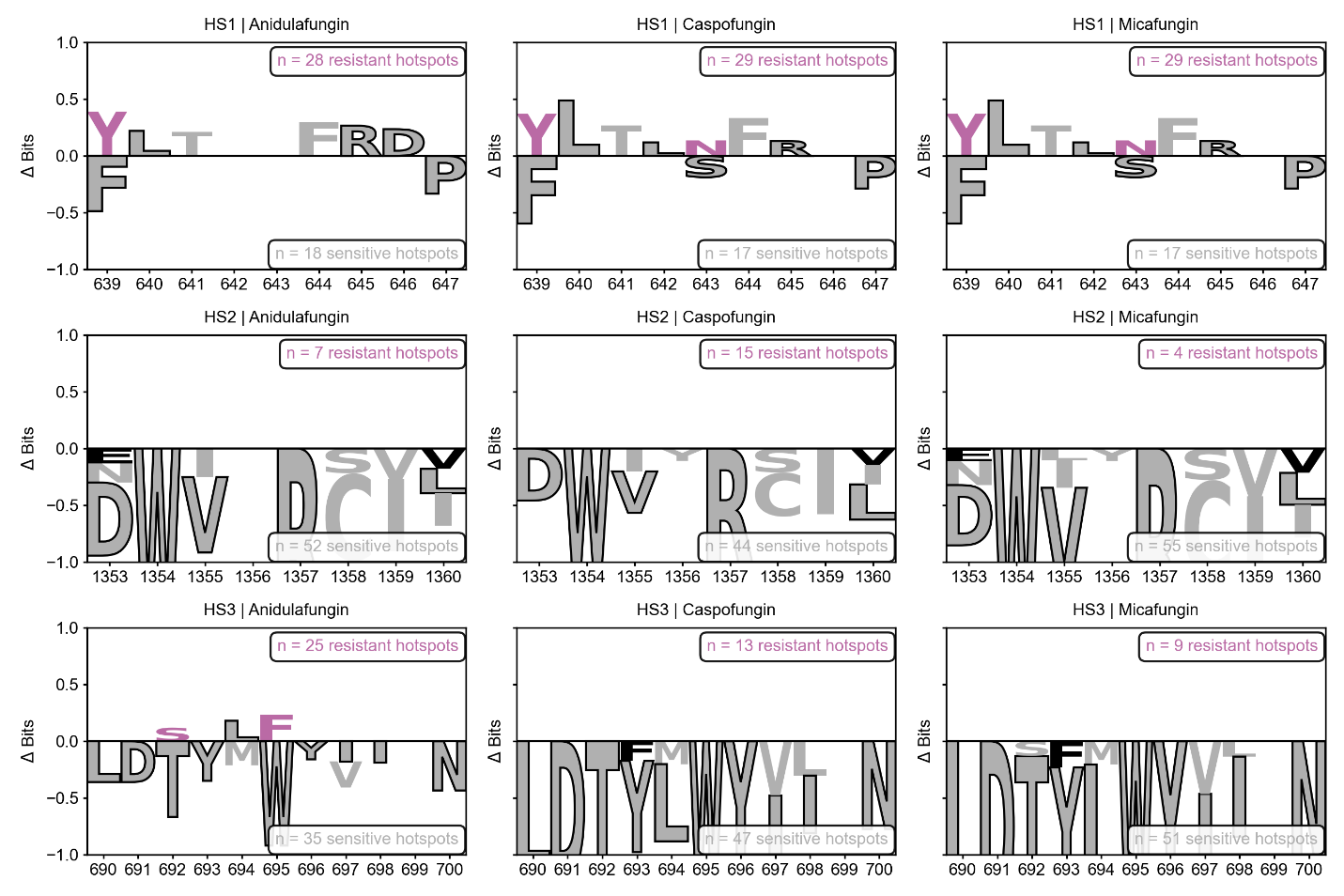

**S11 Figure.** Enrichment analysis of classified homologous hotspot sequences. Enrichment logos for homologous hotspot sequences, classified as resistant (positive) or sensitive (negative) to the indicated echinocandin. Hotspot sequences were retrieved from the multiple sequence alignment used to build the phylogeny tree (S3A Fig). Libraries were built from codon-optimized sequences and spiked with single mutant libraries in the DMS experiment. Selection coefficients for homologous hotspot sequences were then converted to a resistance label using the same classification as for single mutants. The relative occurrence of each amino acid residue at each position in the homologous hotspot sequences was converted to information content measured in bits, where 0 bits represents a random distribution and log2(20) ≈ 4.32 bits represents 100% conservation. The values shown represent the difference in information content (Resistant - Sensitive). Positions with an absolute difference below 0.1 bits were masked to reduce visual clutter due to noise. Consequently, the height of each letter reflects the degree of enrichment and conservation at that position, with positive values indicating residues associated with resistance and negative values indicating residues characteristic of sensitive hotspots. Residues are colored pink if the corresponding substitution (in *S. cerevisiae*) was classified as resistant in the DMS, while residues colored gray correspond to sensitive substitutions. Residues with a black stroke correspond to the wild-type hotspot sequence in *S. cerevisiae*. Logos were generated with Logomaker v0.8.7 (Tareen and Kinney 2020).

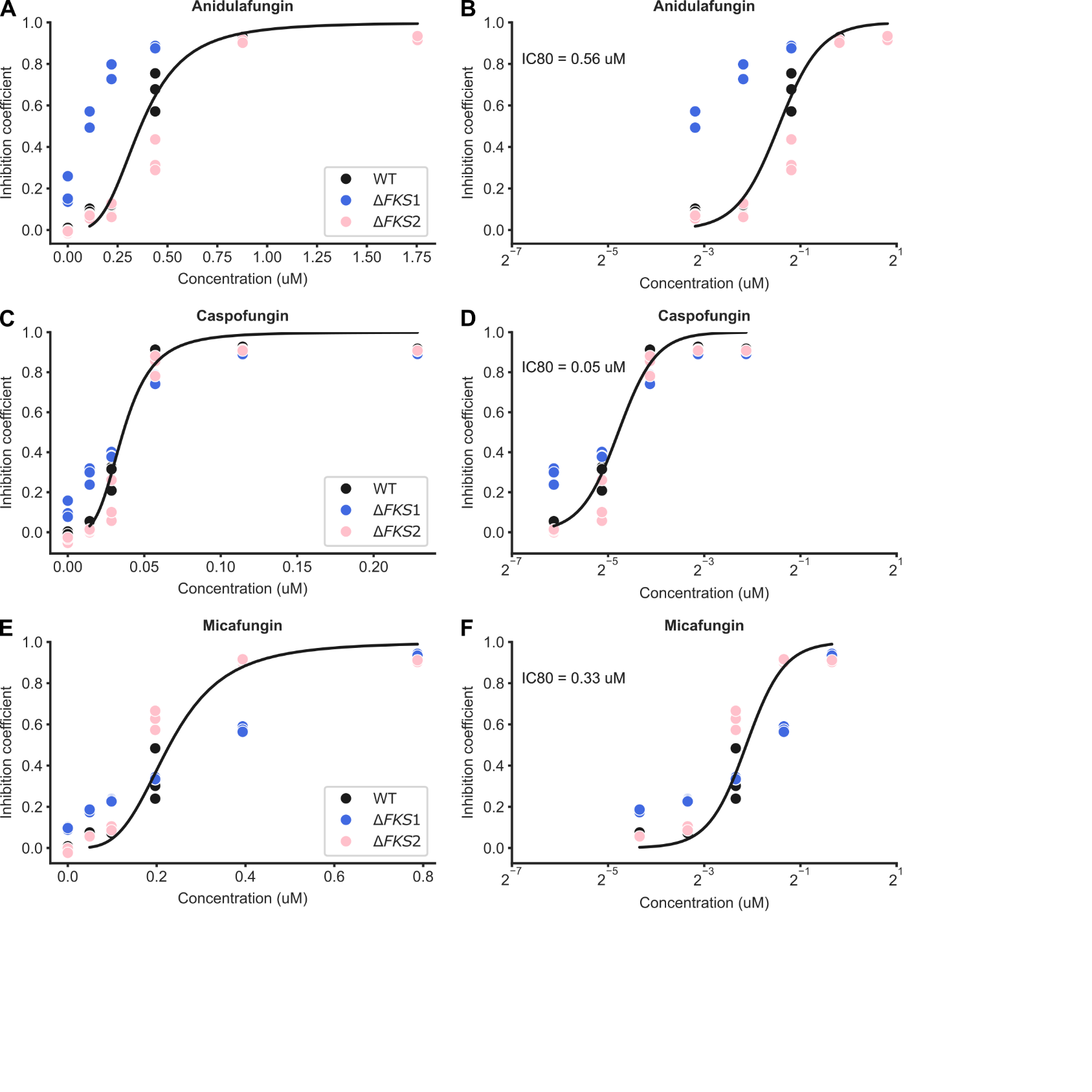

**S12 Figure. Dose-response curves for anidulafungin, caspofungin and micafungin for different yeast genotypes.**

Inhibition curves for the parental strain (BY4741), and isogenic strains in which either the hotspot 1 of *FKS1* or the hotspot 1 of *FKS2* was replaced by a selection marker, in anidulafungin (**A-B**), caspofungin (**C-D**), or micafungin (**E-F**). Concentrations are shown on a linear scale for panels A, C and E or on a log2 scale for panels B, D and F. 0 µM was used to normalize maximum growth rates obtained from biological triplicates. The resulting inhibition coefficients (1 - growth rate) were used to fit the Hill equation for the WT only. The corresponding IC80 (concentrations used for the validation assay) are indicated in panels B, D and F.

**Data S1. (separate file)**

List of single mutations in hotspots 1, 2 and 3 of Fks1 and Fks2 reported in studies that assessed echinocandin resistance. The list was obtained by parsing the FungAMR database (Bédard et al. 2025), which annotates observations with a degree of evidence. All entries corresponding to the present study were filtered out but are available in FungAMR. Degrees of evidence can be positive (resistance) or negative (sensitivity) and include:

1. Mutation created in the same species background and effect on resistance confirmed by measurement
2. Mutation created in the same gene but expressed in another species with potentially non endogenous level of expression
3. Effect of mutation quantified by DMS or similar approach
4. Mutation identified by experimental evolution and high frequency of independent evolution suggests it causes resistance
5. Deletion of the gene causes resistance
6. Overexpression or duplication of the gene causes resistance
7. GWAS (or other population-based association such as QTL or classical genetics): significant association between mutation and resistance in population
8. Mutation identified in a 'natural' (or experimental if no replication as in 4) strain that is resistant but with no specific validation beyond this point; considered as strong candidate mutation because of annotations or other indirect evidence.

For example, “-1” would indicate that the mutation was recreated in the same species background and found to be sensitive to the drug. The best resistance/sensitivity scores reflect the best degree of evidence out of all observations of resistance/sensitivity (e.g. same mutation reported in distinct studies). In the present study, we refer to “high-confidence reports from the literature”: those correspond to entries with codes 1, 2, 3 or 4 (or -1, -2, -3 or -4 for observations of sensitivity).

**Data S2. (separate file)**

Comparison between the phenotypes of single Fks1 hotspot mutants observed here and the ones reported in the literature. S1 Data was filtered to only keep Fks1 mutants for which there was at least one confident report of drug resistance (or sensitivity), i.e. a score between -4 and 4 (see caption of S1 Data for more details on scores) for caspofungin, micafungin or anidulafungin. The best degree of evidence (minimum out of all observations between the absolute values of the best resistance score and the best sensitivity score) was converted to a resistance label (“resistant” or “sensitive”). Columns "Anidulafungin", "Caspofungin" and "Micafungin" indicate if the mutant was classified as resistant or sensitive to the drug in both this assay and in other isolates as previously reported. The value "disagreement" indicates that the phenotype reported here is different from what was reported by others. The discrepancy may be explained by either the quality of the reported data, reflected in the FungAMR scores, and/or the evolutionary distance (see last column) and/or other factors (different media, different drug concentrations, different methods for measurement and/or classification, etc).

**Data S3. (separate file)**

List of homologous Fks hotspots.

**Data S4. (separate file)**

List of strains, plasmids and primers used in this study.
